# HaloTag Ligand and HaloTag Protein engineering for a binary fluorescent turn-on probe

**DOI:** 10.64898/2026.05.15.724826

**Authors:** Blaise Gatin-Fraudet, Ulrich Pabst, Christina H. Olesen, Bianca C. Baciu, Ramona Birke, Sigrid Milles, Johannes Broichhagen

**Affiliations:** Leibniz-Forschungsinstitut für Molekulare Pharmakologie (FMP), 13125 Berlin, Germany

**Keywords:** HaloTag, Protein Engineering, fluorogenicity, microscopy, single molecule imaging

## Abstract

Protein labelling by covalent attachment of a specific substrate to a self-labelling protein tag has become a regular in the life sciences. Herein, we report the design of a two-component labelling system, comprised of a non-fluorescent difluorinated xanthene, called F_2_X, and a HaloTag mutant engineered for targeted reactivity towards F_2_X. Upon primary covalent locking of the ligand at the canonical aspartate residue, two proximal lysine residues located at the protein surface can undergo nucleophilic aromatic substitution with the F_2_X core, building a fluorescent rhodamine via triple-covalent fusion. We used a generalizable *in silico* pipeline for heuristic conformational sampling of covalent protein-ligand complexes to find suitable mutation sites, culminating in the curation of 7 double-lysine HaloTag mutants for targeted *in vitro* testing. Reaction with the best-performing mutant, HTP^L161K_Q165K^, is characterized by full protein mass spectrometry, fluorescence polarization fluorescence lifetime, and fluorescence anisotropy and rationalized by computational modelling. We showcase the system in single molecule microscopy, where obviation of post-labelling purification is a prime advantage when targeting recombinant proteins that may not be expressed in larger quantities, and employ F_2_X in living cells with reduced photobleaching. Lastly, a cell-impermeable version was obtained by means of sulfonation, exclusively targeting extracellularly exposed HTP^KK^ fused to the neuromodulatory G protein-coupled receptor metabotropic glutamate receptor 2.

## INTRODUCTION

The HaloTag Protein, engineered from the *Rhodococcus* dehalogenase DhaA, has become a versatile labelling platform when genetically fused to proteins of interest.^1^ Apart from being covalently reacted with molecules for instance for calcium imaging^2–4^ or FRET based sensors for membrane potential^5,6^, its most often employed application is in fluorescence microscopy.^7^ In line with this, chemical development spurred the quest for bright and stable fluorophores^8^ as well as optimizing its HaloTag Ligands (HTLs)^9^, chloroalkane based substrates. To date, a massive input in rhodamine-based fluorophore optimization, apart from reaching higher brightness, has been addressed to make molecules more fluorogenic, as they exist in two isomers in solution, a non-fluorescent spirolactone and a fluorescent zwitterionic form, which is defined by its equilibrium constant *K*_L-Z_ (**Fig. 1**)^10–12^. Ideally, the dye remains dark in any environment, yet when it binds to HTP, the open form should dominate. Turn-on values of >100-fold have been achieved for carbopyronine or molecular rotors,^12,13^ yet a truly fluorogenic probe would be binary, which in turn would neglect any need for washing to remove unbound ligand.

**Fig. 1.**
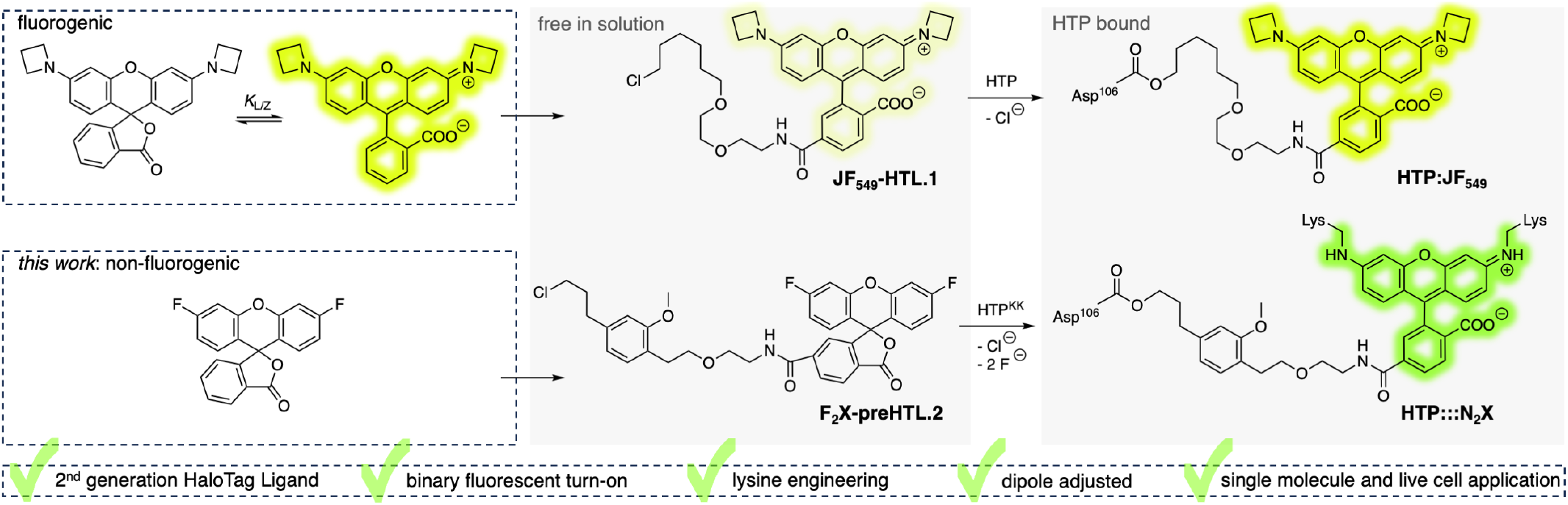
Fluorogenic HaloTag Protein Labelling. Fluorogenic dyes rely on the equilibrium (K_L/Z_) between dark spirolactonic (left) and fluorescent zwitterionic (right) isomers. When free in solution they exhibit dim fluorescence, which is drastically enhanced when bound to a HaloTag Protein (HTP) through covalent linkage via a bio-orthogonal HaloTag Ligand (HTL). Our strategy aims to achieve a binary fluorescent turn-on response enabled by a strictly spirolactonic form. Upon reaction with HTP the difluoroxanthene (F_2_X) stays dark until two nucleophilic aromatic substitutions generated the rhodamine dye solely on the protein surface, ultimately forming three covalent bonds.

In this study, we aimed to tackle this issue from a chemical and a protein engineering side, by 1) designing and synthesizing a dark ligand that can be coupled to HTP, 2) translating 130-year-old chemistry—Meisenheimer-type nucleophilic aromatic substitution reactions—to lysines on the HTP surface to 3) yield a binary fluorescence turn-on system (**Fig. 1**). After designing and synthesizing a difluoro-xanthene (F_2_X) HaloTag ligand, screening a subset of potential lysine substitutions on the HTP surface, and careful characterization and benchmarking using mass spectrometric and bulk and single molecule fluorescent polarization and anisotropy analysis, respectively, we show its application on the single molecule level and in live cell labelling, where additional purification steps post-labelling become unnecessary. Although reaction kinetics are suffering in this first-generation method, we envision that our insights from protein design and reactivity serve as a catalyst for future labelling and imaging perspectives.

## RESULTS

Our first aim was to completely supress fluorogenicity of a dye with a xanthene core, like JaneliaFluor 549 (JF_549_)^14^, to its dark spirolactone form (**Fig. 1**). The equilibrium (*K*_L-Z_) between the fluorescent and dark isomers is affected by several chemical factors, *i*.*e*. the nature of 1) nitrogen substitution that prefers the open form when electrons are pumped into the xanthene core,^15–18^ 2) the nucleophilicity of the chemical group in the 3’-position that has been described to range from carboxylates, alcohols,^12,20^ acyl sulfonamides,^21^ to adapt the dark form, and 3) the installation of charges on the fluorophore core that create a local polar environment to push the open spirolactone form.^22,23^ The dye is therefore in equilibrium partially in the fluorescent form, which may lead to background fluorescence, yet it opens up due to the protein surface of the HaloTag once covalent linked that has been engineered to support and stabilize the zwitterionic form.

Our idea was to replace the *N,N*-bis-alkylated nitrogen groups, which promote electron delocalization (as indicated by typical Hammett constants of σ_p_ = −0.83),^24^ with an electron-pulling moiety that constrains the molecule to its spirolactonic form. Naturally, being both the most electronegative element that the periodic table has to offer and the least sterically demanding unit that could be introduced to the system, fluoride substitution was chosen (σ_p_ = −0.06)^24^. We reasoned that this difluoroxanthene (called F_2_X), being unable to open to the zwitterionic form and therefore lacking complete π-conjugation in the xanthene core, would not be fluorescent.

HaloTag Protein labelling can then be performed when F_2_X is chemically fused to a HaloTag Ligand (HTL), and we chose the second generation, HTL.2 (ref^9^), which has been used in studies to down-titrate the ligand concentration as its affinity towards the HTP orthosteric site has been increased.^9,25^ Although HTL.2 comprises an additional benzoyl group for better labelling, this moiety is encompassed by the lower ring of the F_2_X, and we therefore decided upon F_2_X-preHTL.2. We next opted for a functional group to activate the ligand only when bound to HTP, and thought about amino acid side chains that may serve as nucleophiles to replace the fluoride atom. Lysine may be the ideal candidate, as its side chain carries a primary amine, which in turn would build a fluorophore on site. The drawback would be that lysines on protein surfaces are protonated in the physiological environment (p*K*_a_ ~ 10.4, although within a protein they may become acidic)^26^, however, we were inspired by the fact how nature achieves selectivity, *i*.*e*. by entropically trapping substrates and bringing their “poor” electrophiles close to the nucleophile (beautifully exemplified by HTP itself^1^). Since F_2_X would be connected to HTP by one covalent bond through Asp106, we withered a chance that a lysine may undergo a nucleophilic aromatic substitution to replace a fluoride and yield an aniline, ideally two times.

Therefore, we set out in parallel to synthesize F_2_X-preHTL.2 from a bistriflate xanthene (**1**) to first install the fluoride by means of Buchwald-Hartwig coupling using CsF and [Pd(cinnamyl)Cl]_2_ as a transition metal source and *t*BuBrettPhos as a ligand (ref^27,28^), to be then *t*-butyl ester saponified by TFA in DCM for final installation of the HTL (**Fig. 2A**, upper part, see Supporting Information). Similar chemistry, but with *N*-Boc-ethylamine as a coupling partner and Pd_2_ dba_3_/XantPhos as a catalytic system yielded a dummy compound (denoted as (EtNH)_2_X) that mimics the final reaction product on the HTP (**Fig. 2A**, lower part). We next checked the UV/Vis spectra of free F_2_X-6-COOH and (EtNH)_2_X-6-COOH, and, as expected, observed maximal absorption in the UV region and around 520 nm, respectively (**Fig. 2B**), proving the full closure to the spirolactone form for F_2_X-6-COOH. The latter is further supported by ^1^H and ^19^F NMR spectroscopy that show only one species (see Supporting Information). Labelling of the ligand was performed by incubation with recombinantly expressed HTP and subsequent measurement of full protein mass spectrometry, which gave a complete and mono-covalent labelling reaction (**Fig. 2C**).^25^ When recovering these labelled samples and running emission scans with λ_Ex_ = 500 nm, we were only observing noise for the F_2_X-labelled, but clear fluorescence of the (EtNH)_2_X-labelled HTP, giving confidence that a binary turn-on might be feasible (**Fig. 2D**).

**Fig. 2.**
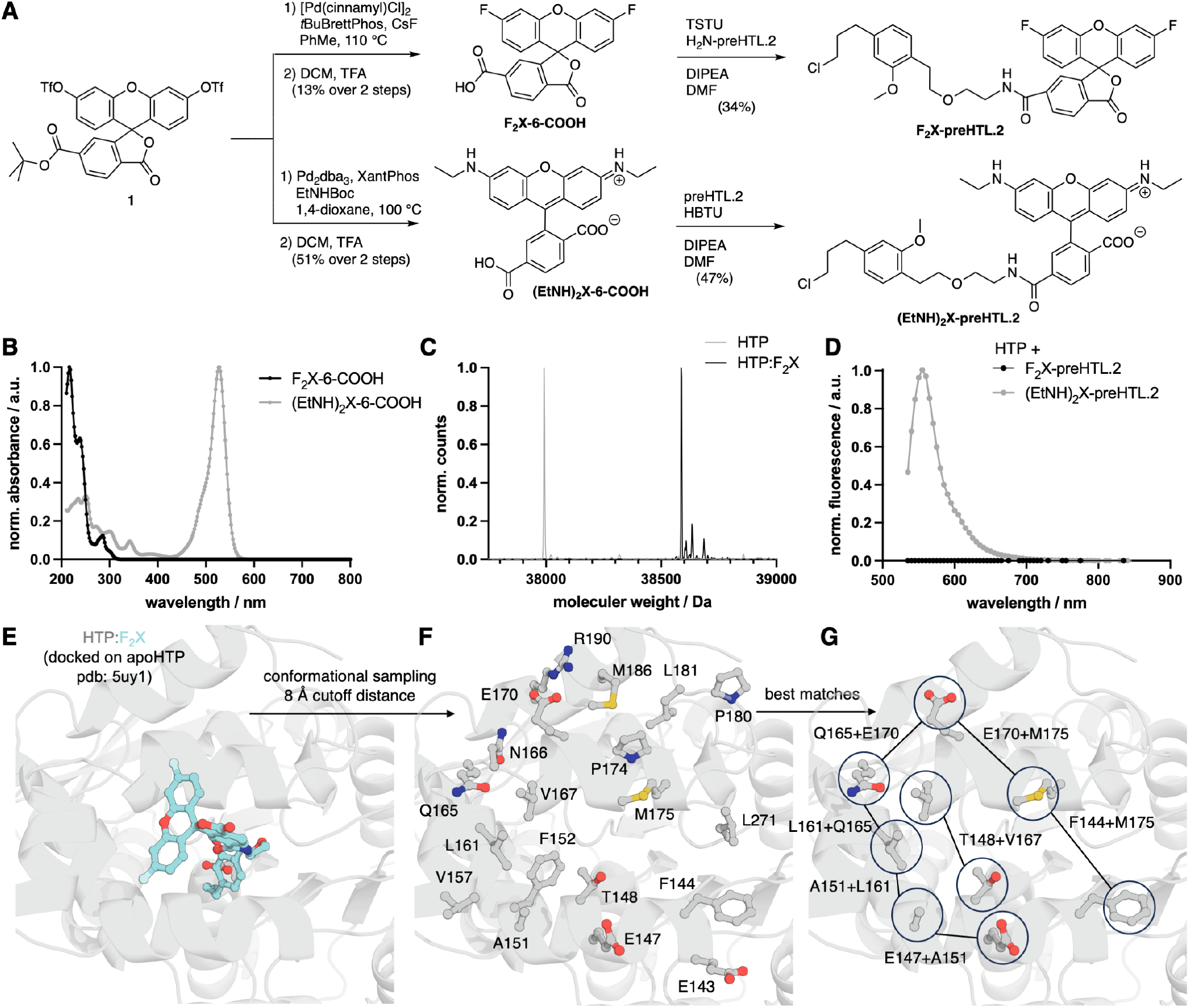
Synthesis and Protein Design of the binary F_2_X label system. **A)** Synthesis of F_2_X-preHTL.2 (left) and (EtNH)_2_X-preHTL.2 (right). **B)** Absorbance profiles of F_2_X-6-COOH and (EtNH)_2_X-6-COOH. **C)** *In vitro* protein labelling of apo-HTP by F_2_X-preHTL.2 with mono-covalent binding confirmed by intact protein mass spectrometry after 2h. **D)** Emission profiles of HTP:F_2_X and HTP:(EtNH)_2_X. **E)** Computational model of F_2_X bound to HTP active (based on PDB: 5uy1). **F)** Conformational sampling of the xanthene core highlights all amino acids within an 8 Å cutoff distance. **G)** Representation of all double-lysine HTP combinations that are envisioned as best matches for aromatic substitutions on F_2_X.

Simultaneously, in order to effectively activate fluorescence of the supplied F_2_X substrate upon conjugation to HTP, we had to provide two lysine residues on the protein surface that are optimally oriented to attack the fluoride-substituted carbon atoms on the xanthene moiety on both sides. This was intended for promoting formation of the triple-covalent complex, leading to full π-conjugation across the xanthene core and thus activating fluorescence of the fully assembled conjugate. Given the known overall stability of HTP, as well as its specific tolerance for extensive lysine mutations on the protein surface shown by Koßmann et al.,^29^ we anticipated pairwise lysine mutations to not interfere with the overall structure, dynamics, and reactivity of the protein.

However, as we intended to promote the lysine substitutions by prolonged, forced exhibition, strategic positioning of the mutated residues was considered crucial. Since published crystal structures of HTP:dye conjugates regularly indicate a certain conformational flexibility of the ligand within the cavity (*e*.*g*., PDB-6U32),^30^ we used computational tools to identify potential candidates for pairwise mutation and subsequent *in vitro* testing (for details, see Supporting Information).

First, we covalently docked F_2_X-preHTL.2 into the HTP cavity (as obtained from the apo-HTP crystal structure in PDB-5UY1),^31^ where the covalent bond was formed with the carboxylate of Asp106 (**Fig. 2E**). The resulting complex, HTP:F_2_X, was then subjected to explicit-solvent molecular dynamics simulation to account for the flexibility of both the tethered ligand and the surrounding protein side chains.^32^ After equilibration, a production trajectory was analyzed by reducing each residue in every frame to the Cartesian centroid of its side-chain heavy atoms. For each frame, distances were calculated between all side-chain centroids and the two fluorine-bearing carbon atoms of the xanthene core, being the respective electrophiles. Residues were then ranked by two complementary criteria: the fraction of trajectory frames in which the side-chain centroid was located within 8.0 Å of either electrophilic carbon, and a cutoff-free inverse-distance score that weights persistent close approaches over the full trajectory (**Supporting Fig. S1**). This MD-based sampling identified a focused set of surface-accessible residues proximal to the reactive xanthene positions, from which pairwise lysine mutation candidates were selected for experimental testing after visual inspection of the highest-ranking sites (**Fig. 2F,G**).

To put things together, we next mutated the seven double-lysine versions in a plasmid for recombinant expression (**Table 1**), and produced all of these in good yields and high purity from *E. coli* BL21 (DE3) cells by Ni-NTA affinity chromatography (**Supporting Fig. S2** and see Supporting Information). Next, we performed the critical experiment and incubated the mutants (1 µM) with F_2_X-preHTL.2 in PBS (pH = 7.65) over night in a 96 well plate, and recorded emission spectra the next day. Gratifyingly, three mutants showed a strong increase in fluorescence emission at the expected maximal wavelength λ_Em_ = 550 nm (λ_Exc_ = 500 nm), with mutant HTP^L161K_Q165K^ showing the highest intensity (**Table 1** and **Fig. 3A**).

**Table 1.** HTP mutations and characterization by full protein mass spectrometry and relative Fluorescent Turn-on with F_2_X-preHTL.2. (1 µM protein, and 2.5 µM F_2_X-preHTL.2, incubation for 16 h at 37 °C).

| HTP mutation | expected mass (kDa) | found mass (kDa) | Rel. fluorescent turn-on vs. BSA (%) |
| --- | --- | --- | --- |
| none | 37990 | 37992 | 0.4 ± 0.2 |
| L161K, Q165K | 38005 | 38005 | 100 ± 0.7 |
| Q165K, E170K | 38011 | 38010 | 58.2 ± 0.3 |
| E170K, M175K | 37985 | 37987 | 4.4 ± 0.4 |
| A151K, E147K | 38045 | 38047 | 2.3 ± 0.3 |
| A151K, L161K | 38061 | 38062 | 36.2 ± 0.7 |
| F144K, M175K | 37968 | 37968 | 6.3 ± 0.4 |
| T148K, V167K | 37873 | 37872 | 54.7 ± 0.7 |

**Fig. 3.**
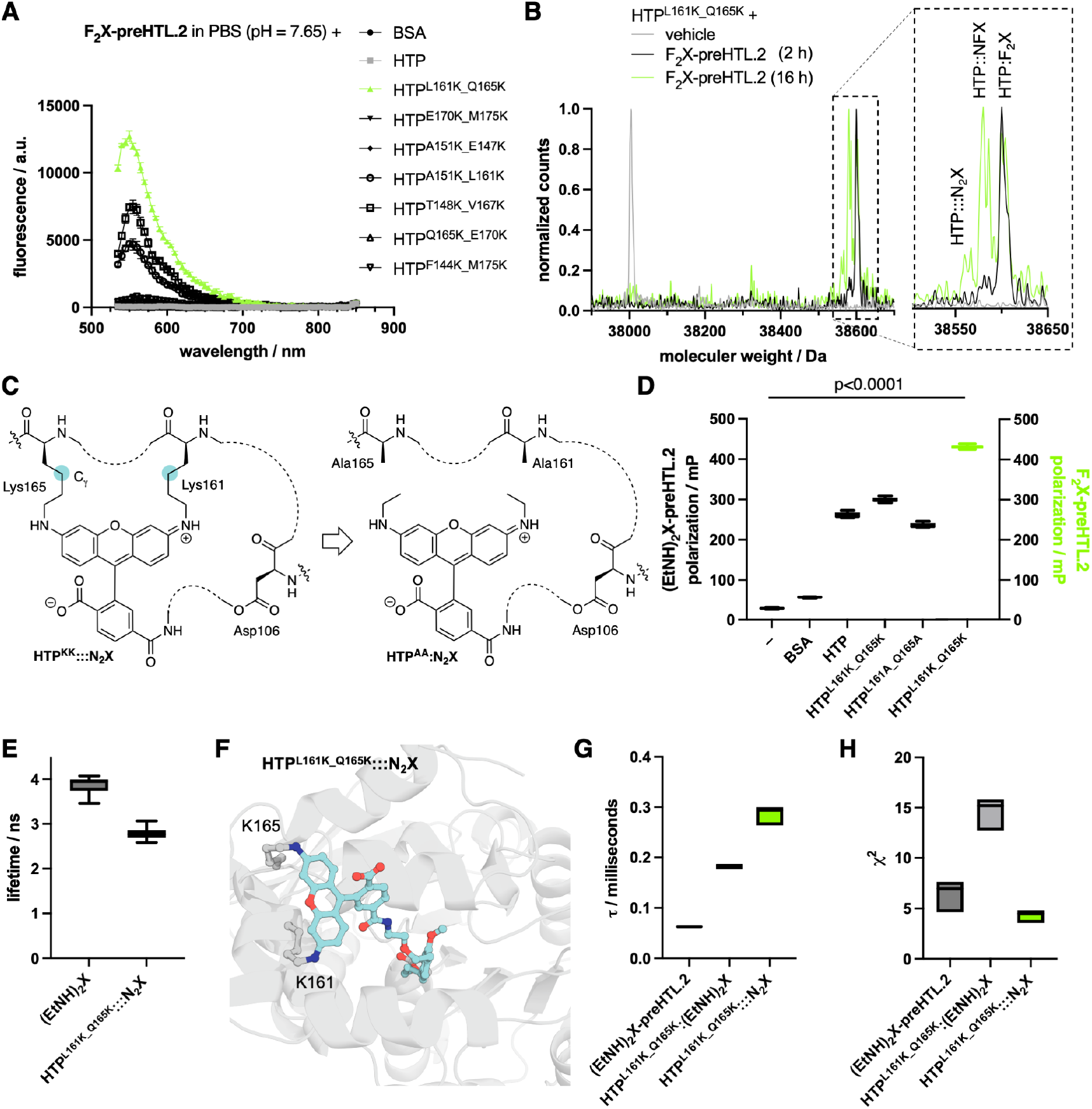
Characterization and Photophysical properties of the binary label system. **A)** Raw fluorescence of 2.5 µM of F_2_X-preHTL.2 and BSA, HTP and all the HTP mutants in PBS (pH = 7.65) after 16h incubation at 37 °C. (n = 3). **B)** Intact protein mass spectrometry of apoHTP^L161K_Q165K^ or with F_2_X-preHTL.2 after 2h and after 16h. **C)** A less restricted dye as a control versus triple bound F_2_X is achieved by using (EtNH)_2_X and a double lysine to alanine mutation, formally cutting the C_γ_. **D)** Fluorescence Polarization of (EtNH)_2_X-preHTL.2 alone, or in presence of BSA, HTP, HTP^L161K_Q165K^, HTP^L161A_Q165A^ and F_2_X-preHTL.2 bound to HTP^L161K_Q165K^ after 16h incubation at 37 °C. Box-and-whisker min-max, n=8. **E)** Fluorescence lifetime of F_2_X-preHTL.2 and (EtNH)_2_X-preHTL.2 after 16h incubation at 37 °C with HTP^L161K_Q165K^. Box-and-whisker min-max, n=39 ((EtNH)_2_X) and n=33 (F_2_X). **F)** Modelling of the HTP^L161K_Q165K^ active site bound to F_2_X-preHTL.2 forming HTP^L161K_Q165K^:::N_2_X. **G)** Diffusion times extracted from fits of fluorescence correlation spectroscopy (FCS) curves, where a single diffusion time was imposed. Samples recorded were (EtNH)_2_X-preHTL.2, bound to HTP^L161_Q165K^ and HTP^L161_Q165K^:::N_2_X. Floating bars min-max. n=3. **G)** Corresponding χ^2^ values for (F), assessing the quality of the fit with a single diffusion time. Floating bars min-max. n=3.

Owing to the basic nature of lysine’s side chain amines that are protonated, we repeated these incubations, yet at non-physiological pH = 8.65 and 9.15, resulting in a 3.6- and 3.9-fold fluorescence emission when compared to pH 7.65 (**Supporting Fig. S3**). Under both conditions we observed larger signal in fluorescence emission and a slight bathochromic shift, more pronounced for pH = 9.15, indicating how sensitive the microenvironment is for both the reaction and fundamental photophysical properties. Equally important, incubation with bovine serum albumin or the native HTP did not lead to any fluorescence (**Fig. 3A**). As such, we took the same sample of HTP^L161K_Q165K^ and performed full protein mass spectrometry, finding all species: mono-reacted HTP:F_2_X, doubly reacted HTP::NFX and triple reacted HTP:::N_2_X, with a mass shift of 20 Da, as expected when considering that formally hydrogen fluoride is liberated (**Fig. 3B**). We also investigated the size of the HaloTag Ligand, and testing HTL.1, and HTL.2, we confirmed that F2X-preHTL.2 exhibited the highest turn-on after 16 h of incubation (**Supporting Scheme S1** and **Supporting Fig. S4**). To better assess the efficiency of the system, we calculated the ratio of fluorescence intensity after incubating F_2_X-preHTL.2 and (EtNH)_2_X-preHTL.2 with HTP^L161K_Q165K^ (for easier reference now referred to as HTP^KK^) for 16 h at 37 °C. This analysis suggests that less than 5% of F_2_X-preHTL.2 was converted into the fluorescent product (**Supporting Fig. S5**). While this is considering photophysical properties to be identical (which they are not), we further quantified full reaction yield when calculating the area under curve from full protein mass spectrometry (**Fig. 3B**), under the assumption that ionizability does not change, where we obtain a triple covalent linkage yield of 16% (**Supporting Fig. S5**). As such, we attribute a triple covalent linkage to be yielded 210% after 16 hours.

Since the dipole is adjusted by three non-co-linear attachment points define a plane, we were interested if this is reflected in fluorescence polarization. In order to mimic the steric demand that an *N*-ethyl group brings, we also cloned, expressed, purified and labelled HTP^L161A_Q165A^ (for easier reference now referred to as HTP^AA^) with (EtNH)_2_X-preHTL.2, which impersonates HTP:::N_2_X by formally cutting out the Cγ atom from the lysine side (*i*.*e*. 4 carbons in the side chain vs. 1 carbon from alanine plus 2 carbons from *N*-Et, **Fig. 3C**). Next, we recorded fluorescence polarization of free (EtNH)_2_X-preHTL.2 in PBS buffer and BSA excited with polarized light (~50–60 mP), and when bound to HTP and HTP^KK^ (~260-300 mP) (**Fig. 3D, Table 2**), indicating lost degrees of freedom, and showcasipolarization was observedng how subtle changes in the environment influence fluorophore tumbling.

**Table 2.** Fluorescence Polarization of HTP mutants. n = 10, nd = not determinable.

| Polarization / mP | (EtNH)<sub>2</sub>X-preHTL.2 | F<sub>2</sub>X-preHTL.2 |
| --- | --- | --- |
| Vehicle | 28 ± 2 | nd |
| BSA | 56 ± 2 | nd |
| HTP | 261 ± 7 | nd |
| HTPKK | 299 ± 5 | 431 ± 4 |
| HTP<sup>AA</sup> | 235 ± 6 | nd |

Similarly, for the HTP^AA^ mutant labelled with (EtNH)_2_X-preHTL, fluorescence polarization was observed at ~230 mP (**Fig. 3D**). Importantly, HTP:::N_2_X showed a strongly increased fluorescence polarization at ~430 mP, demonstrating the entropic surge of triple covalent modification (**Fig. 3D, Table 2**).

To further characterize our system, we obtained fluorescence lifetimes of HTP^L161K_Q165K^:::N_2_X and (EtNH)_2_X-6-COOH, in which the conformationally locked version showed faster return to the electronic ground state, therefore minimizing its lifetime (τ = 3.6 ns vs. τ = 2.1 ns) (**Fig. 3E**). To provide a visual representation of the proposed HTP:::N_2_X structure, we generated an illustrative structural model of the triple-covalent complex (**Fig. 3F**). Starting from the HTP scaffold, the L161K/Q165K mutations were introduced in ChimeraX using Dunbrack rotamers,^33,34^ and F_2_X-preHTL.2 was positioned in the binding cavity through covalent docking to Asp106 (see Supporting Information).

The model was then manually converted into the expected triple-covalent product by opening the spirolactone, replacing the aryl fluorides with lysine linkages, and adjusting the corresponding bond orders. The ligand and nearby cavity residues within 8.0 Å were subjected to backbone-constrained MMFF94 energy minimization in Avogadro,^35,36^ followed by reintegration into the full protein model and short molecular dynamics relaxation. This model was used solely for visualization of a chemically plausible product and should therefore be interpreted as a representative model rather than a predicted minimum-energy structure.

We also calculated fluorescence correlation curves of the different HTL variants. Even though, technically, all protein variants have the same size and should thus diffuse through the confocal volume with a similar speed, we observed faster diffusion times for all protein variants compared to HTP^KK^:::N_2_X (**Fig. 3G**), stemming from contamination by the fluorescent unbound dye (used two-fold excess). Indeed, a single diffusion time did not accurately describe the fluorescence correlation spectroscopy (FCS) curves (see Supporting Information) in those cases, also exemplified by a large fitting error if a single diffusion time was used (**Fig. 3H**). This is different for the HTP^KK^ variant, where, strictly, only the bound dye is fluorescent. The FCS curves of HTP^KK^:::N_2_X were very well described with a single diffusion time and showed a very small fitting error (**Fig. 3G, H**), enabling direct use without further purification.

We next tested the behaviour of F_2_X in a cellular environment, and introduced a double alanine and a double lysine mutant on positions 161 and 165 on an HTP-SNAP-NLS construct^12^ that localizes to the nucleus (NLS = nuclear localization sequence). After transfecting HEK293T cells with HTP^AA^-SNAP-NLS and applying (EtNH)_2_X-preHTL.2 as well as BG-SiR-d12 (for SNAP-tag labelling)^37^, we observed signals from the nucleus in both channels by confocal microscopy (**Fig. 4A**). We likewise observed no signal when F_2_X-preHTL.2 was used instead in an overnight incubation (**Fig. 4B**). However, when we transfected the HTP^KK^-SNAP-NLS mutant and labeled with F_2_X-preHTL.2, we were able to detect clear signals from the nucleus. We then recorded 40 frames on the HTP:::N_2_X and HTP:(EtNH)_2_X samples and integrated the fluorescence density of each image before normalizing (**Fig. 4C**). Mono-exponential fitting of the mean values gave a 95% confidence interval for t_1/2_ = 5.73–7.76 frames (plateauing at 0.84) for HTP:::N_2_X with reduced bleaching compared to HTP:(EtNH)_2_X (95% confidence interval for t_1/2_ = 10.7–12.2 frames; plateauing at 0.47), accounting for its improved photostability.

**Fig. 4.**
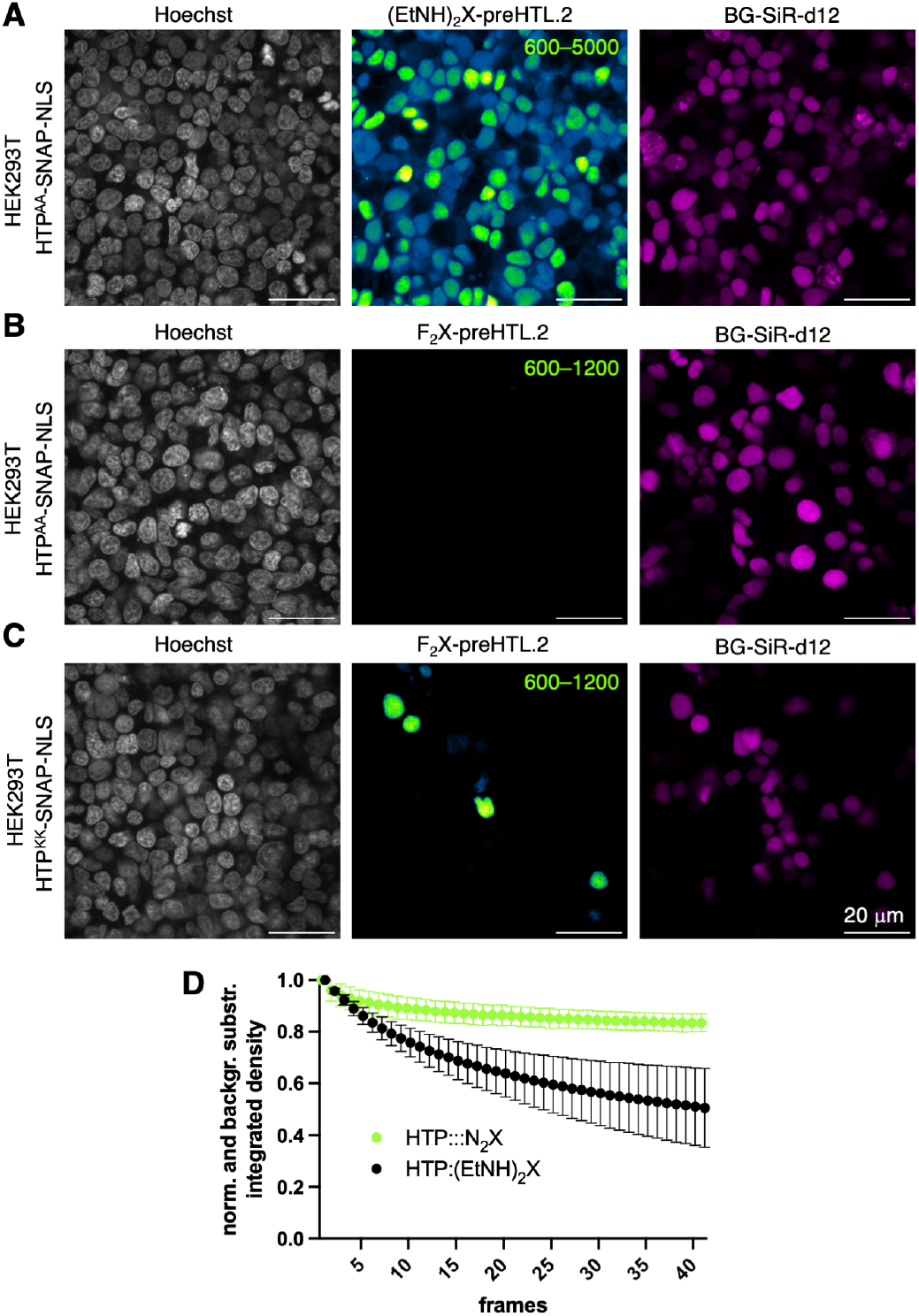
Live cell imaging of F_2_X-preHTL.2 in transfected HE293T cells. **A** and **B)** Confocal images of HEK293T cells expressing HTP^AA^-SNAP-NLS labelled with BG-SiR-d12 and (EtNH)_2_X-preHTL.2 (2.5 µM, A) or F_2_X-preHTL.2 (2.5 µM, B). **C)** Confocal images of HEK293T cells expressing HTP^KK^-SNAP-NLS, HTP labelled with F_2_X-preHTL.2 (2.5 µM). Imaging was performed for all conditions in N = 3 preparations with n = 5 images per condition. SNAP labelled with BG-SiR-d_12_. For all images, scale bar = 20 μm. **D)** Bleaching curves of HTP:::N_2_X and HTP:(EtNH)_2_X of the images from (A, B). mean±SD, N = 3. n = 3-5.

To further expand the repertoire for F_2_X labelling, we applied our recently described ‘shuttle’ approach,^25^ by reacting F_2_X-preHTL.2 with a strong base (NaH) and 1,3-propylene sultone to install a negatively charged sulfonate on the HTL, which renders the resulting F_2_X-SHTL.2 impermeable towards the plasmalemma (**Fig. 5A**). Intended to exclusively stain cell surface proteins from the extracellular side, where G protein-coupled receptors like the neuromodulatory class C metabotropic glutamate receptor 2 (mGluR2) exert their physiological function, we transfected HEK293T cells with a plasmid encoding SNAP-HTP^KK^-mGluR2. Indeed, after incubation overnight with impermeable SNAP-tag ligand BG-Sulfo646 (ref^22^) and F_2_X-preHTL.2, staining was predominant in the far-red channel (**Fig. 5B**). In case for F_2_X-SHTL.2, good overlap between both dyes can be observed on the cell surface, as well as partly internalized receptor pools. The difference in labelling efficiency most likely stems from the increased solubility of F_2_X-SHTL.2, making more ligand available for the reaction with HTP^KK^.

**Fig. 5.**
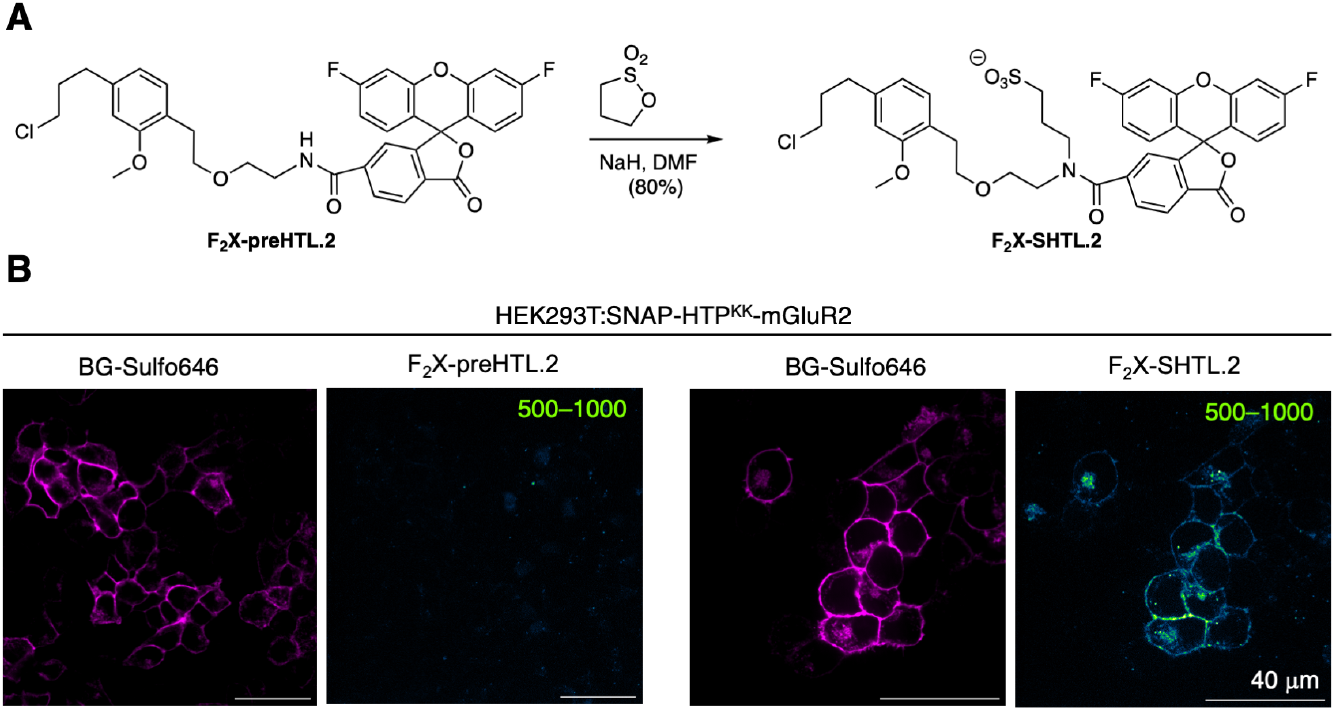
Impermeabilizing F_2_X-preHTL.2 using the ‘SHTL’ approach for cell surface receptor imaging. **A)** Impermeabilizing F_2_X-preHTL.2 via sulfonation, yielding F_2_X-SHTL.2. **B)** Confocal images of live HEK293T cells expressing SNAP-HTP^KK^-mGluR2 treated with impermeable BG-Sulfo646 and F_2_X-preHTL.2 (2.5 µM, left) or F_2_X-SHTL.2 (2.5 µM, right). N = 3, n = 3-5.

## DISCUSSION

Lysines are the prototypical nucleophiles for covalent labelling^38^ and have been–in a more or less specific manner–reacted with acyl halides,^39,40^ NHS esters^41^, isothiocyanates^42,43^, sulfonyl fluorides^44^, or SuFEx^45^, and are in biology^46^ most prominently addressed by acylation and alkylation (*e.g*. for epigenetic regulation), ubiquitination (for degradation) or condensation (*e.g*. with retinal to form rhodopsin). Furthermore, lysines are being selectively addressed, most notably by covalent warheads, most prominently since the late 19^th^ century by acetylsalicylic acid (ASS, Aspirin), and more recently targeting kinases to bind and block the active site,^47,48^ or by *N*-acyl-*N*-alkyl/*N*-acyl-*N*-aryl sulfonamide (NASA/ArNASA)^49,50^, which depicts an affinity labelling approach by pharmacologically directing a reactive group onto a protein surface. However, using aryl fluorides have to date not been described in that regard, as they are mainly used to target cysteines, usually as perfluorinated systems (*e.g*. π-clamp^51^, oxazolinium salts^52^). We showcase such a reaction specifically on a protein surface by not only forming a covalent bound to keep a molecule on site of a protein of interest, but also to introduce direct functionality by converting a difluoro xanthene to a bright rhodamine fluorophore. This is mediated by the HaloTag Protein through a fast capture of the substrate, and by computer guided protein engineering.

Indeed, the HTP has experienced many rounds of optimization since its introduction in 2008.^53^ For instance, labelling kinetics have been boosted from *k*_obs_ = 2.7 × 10^6^ M^−1^.s^−1^ to 1.9 × 10^7^ M^−1^.s^−1^ using tetramethyl rhodamine (TMR).^54^ Furthermore, the surface close to the entry channel has been bulked up with positive charges, leading to the HOB-tag^29^ for enhanced labelling with negatively charged nucleotides and amino acids have been mutated to yield control over fluorescent lifetimes of bound fluorophores.^55,56^ In addition to this, split version have been reported for improving knock-in strategies^57^ and sensor design,^1,58,59^ the latter of which has been extensively probes to record action potential firing optically and beyond.^6,60^ With its main application in imaging by means of protein fusions, we add to the portfolio by introducing two lysines on the surface that are able to react in a nucleophilic aromatic substitution with a difluorinated xanthene (F_2_X) to yield a binary turn-on probe for single molecule spectroscopy and cellular imaging. While the covalent bond formation *via* aspartate106 in the HTP is diffusion controlled as it sits deeply buried within the protein, the second and third reactions are orders of magnitude slower. Although this is obviously expected, we aimed to address this drawback, and found that reactions occur faster at higher buffer pH, and the use of preHTL.2 leads to better results compared to HTL.1 or HTL.2, when delivering F_2_X. We hypothesize that the longer linker of HTL.2 and the higher flexibility of the chloroalkane from HTL.1 allow more flexibility (see **Supporting Scheme 1**), and thereby minimize the ability of the ε-amine of introduced lysines to react. On the flip side, the non-existing fluorescence background stemming from unreacted F_2_X makes this method a versatile tool, in particular for imaging experiments on the single molecule level where highly dilute concentrations are being used.^61^ In a similar vein, locking the fluorophore on three non-colinear points that define a plane, the (transition) dipole of the resulting N_2_X is highly restricted, as can be observed by fluorescence polarization. Since most modern imaging techniques are coming closer to elucidate structures and orientations in structural biology, for instance using MINFLUX that resolves a singularity down to the nanometer level (*i.e*. the size of the fluorophore),^62,63^ or labelling and expansion technologies, such as ONE,^64^ our reported F_2_X strategy may—although a linker error is still in place by the HTP^65^ — contribute to these developments. Furthermore, a more restricted transition dipole may be explored in Förster Resonance Energy Transfer (FRET) between a donor and acceptor molecule, whose efficiency is not only dependent on the distance and overlap integral, but also on the relative orientations. Indeed, obtaining control over the orientation factor κ^2^, one can envision the measurement of energy transfer with prolonged Förster radii. Generally, κ^2^ = 0.66 for isotropic molecules in solution, however, κ^2^ may theoretically adopt values between 0 to 4. As such, when donor and acceptor molecules remain restricted in their anisotropy, an efficiency boost of 35% may be obtained, and with Förster radii nowadays spanning 28 nm, this would result in respective distances of >10 nm. For all the described endeavours, more engineering needs to be undertaken, for instance by 1) adding a second domain on the HTP surface that positions the respective lysines in a more efficient way for nucleophilic attack, 2) further engineering of the dye ‘visible’ HTP surface for more efficient pre-arrangement, or 3) by mutating in histidines and arginines to create a more basic environment for lysine deprotonation (the latter of which is supported by our experiments under more basic conditions, *vide supra*). In order to further orient and lock dipoles, HTP may be inserted into protein of interest *via* a circular permutation (and not as usually *via* its *N*- or *C*-terminus),^66^ restricting its degrees of freedom, and, naturally, the method needs a dye pair as a donor/acceptor.

From a chemical engineering point, F_2_X may become more electron poor by means of electron withdrawing substituents, like the classical example by Fields in the 1970, in which a sulfite became a leaving group by means of tri-nitro aryl substitution.^67^ Similarly, heteroarenes are able to converted F_2_X into a more reactive electrophile,^68^ however, for both cases with potential downsides on its stability and photophysics. Regarding the leaving group, fluoride can be considered ideal due to its high electronegativity and smaller size, compared to its halogen counterparts in the periodic table^69^, and its position is restricted due to the building of a rhodamine. Other colors may be obtained by replacing the central oxygen in F_2_X with *iso*-propylene or dimethyl silyl, to obtain carbopyronines and silicon rhodamines, and generally, scaffolds of fluorophores that bear even one *N*-alkyl group (*e.g*. coumarin, NBD) should in principle be amenable to our approach. Such optimizations are being explored in our laboratories.

## SUMMARY

We expanded protein labelling tools using self-labelling tags by introducing an unprecedented approach: aromatic substitution by highly specific natural nucleophiles on a fluorinated derivative. Through rational design, we developed a non-fluorescent molecule, F_2_X, and engineered seven double-lysine HaloTag Protein mutants *via in silico* conformational analysis. This enabled the identification of a particular double mutant, positions L161 and Q165, which performs best as a binary fluorescent turn-on system, characterized by several techniques, *i.e*. protein mass spectrometry, fluorescence polarization, fluorescence lifetime, and computational modelling. We further demonstrated its applicability in single-molecule microscopy, eliminating the need for post-labelling purification; a critical advantage for recombinant proteins expressed in limited quantities. Additionally, we employed F_2_X in live cells, showcasing reduced photobleaching *vs*. its rhodamine stablemate, and, finally, report a cell-impermeable derivative to selectively label extracellularly exposed HTP^KK^ fused to neuromodulatory G protein-coupled receptor mGluR2 that is crucial in neurotransmission^70^ and involved in pathological states, such asdementia and anxiety.^71–73^ With only two point mutations needed for virtually turning any HTP-fusion into an F_2_X reactive HaloTag, we envision this system—and its successors—as a easy to implement and powerful strategy for protein labelling, spectroscopy and imaging.

## ASSOCIATED CONTENT

### Supporting Information

Chemical synthesis and characterization, computational modelling, measurements of photophysical parameters, and procedures in cell culture, molecular biology and imaging are reported in the Supporting Information.

## AUTHOR INFORMATION

### Author Contributions

Conceptualization and Methodology: JB; Formal analysis and investigation: BG-F, UP, CHO, BCB, RB, SM and JB; Writing –Original Draft: BG-F, UP and JB; Reviewing and Editing: BG-F, UP, CHO, BCB, RB, SM and JB; Visualization: BG-F, UP and JB; Supervision: SM, JB; Funding Acquisition: SM, JB. All authors have given approval to the final version of the manuscript.

## ACKNOWLEDGEMENTS

We are grateful to Leibniz-FMP for support by means of an Integrated Project (SM and JB).

---

Authors are required to submit a graphic entry for the Table of Contents (TOC)

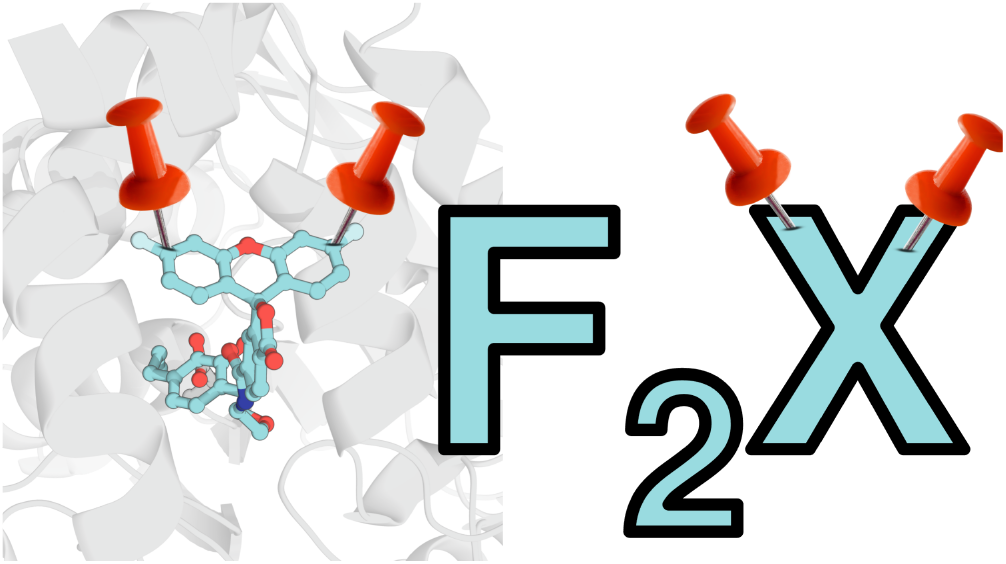

