## Supporting Information for "HaloTag Ligand and HaloTag Protein engineering for a binary fluorescent turn-on probe"

### Supplementary Figures

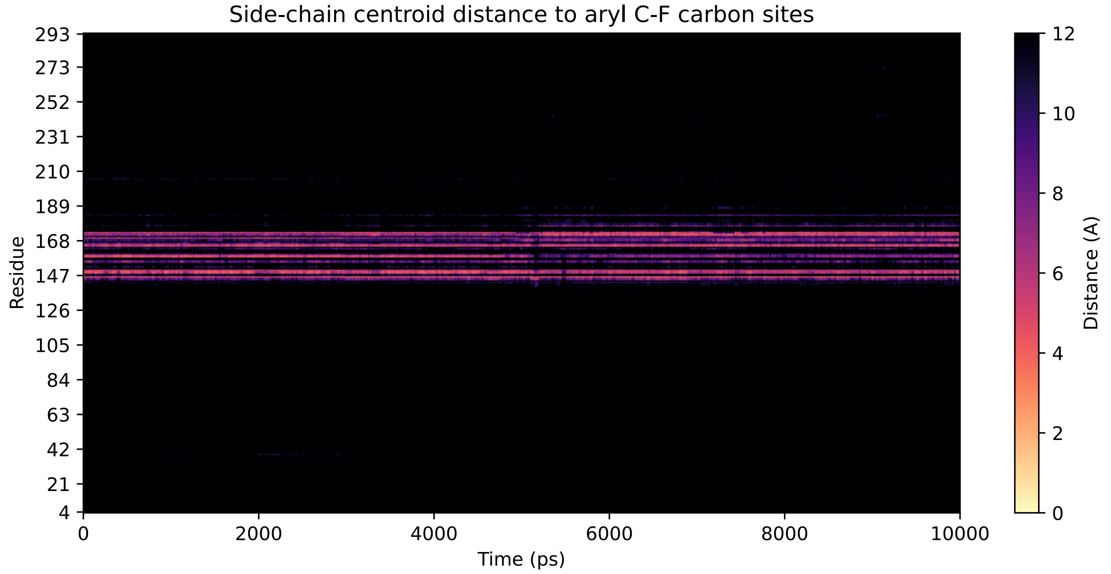

**Supplementary Fig. 1: Conformational Proximity Sampling.** The covalently docked HTP:$F_2X$ structure was equilibrated and simulated using molecular dynamics at 300 K in TIP3P water. For all frames in the 10 ns production trajectory, distances between side-chain heavy atom centroids and the electrophilic carbon atoms on the $F_2X$ core are calculated and shown as the combined minimum distance of each residual centroid to either of the electrophilic carbon atoms.

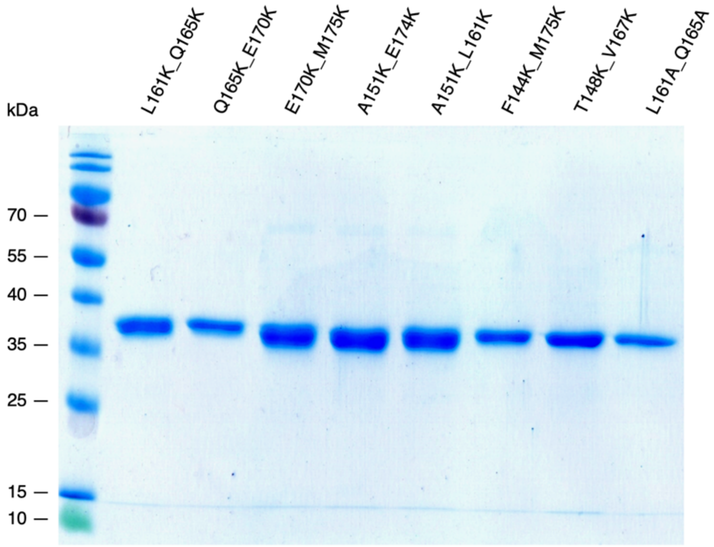

**Supplementary Fig. 2: SDS-PAGE on HTP mutants.**

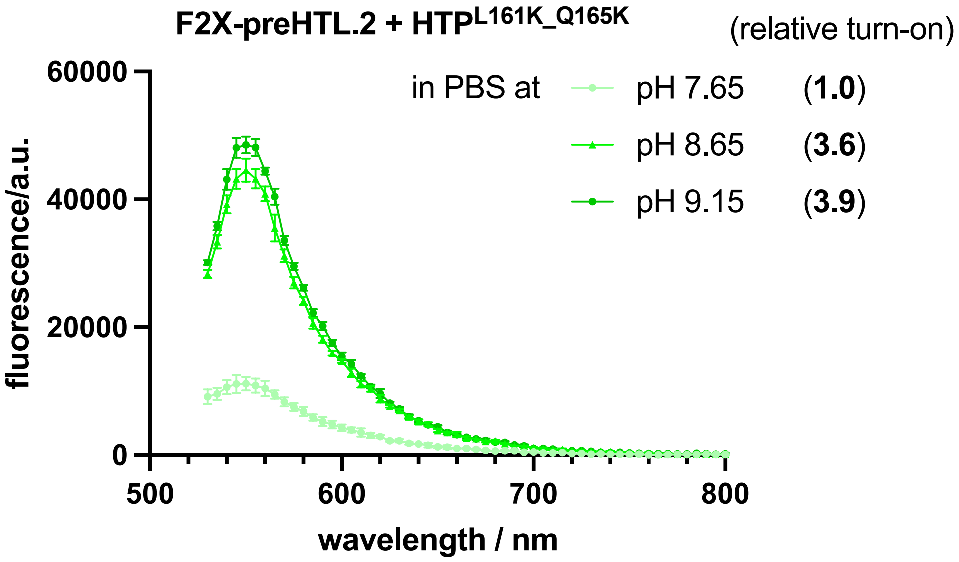

**Supplementary Fig. 3: Influence of pH on the binary system.** Raw fluorescence of 2.5 µM of $F_2$X-preHTL.2 and HTP$^{L161K_Q165K}$ mutant in PBS after 16 h incubation at 37 °C, at pH = 7.65, 8.65 and 9.15 (n = 3). AUC [Total Area (normalized): pH 7.65: 780806 (26%), pH 8.65: 2773202 (92%), pH 9.15: 3012665 (100%)].

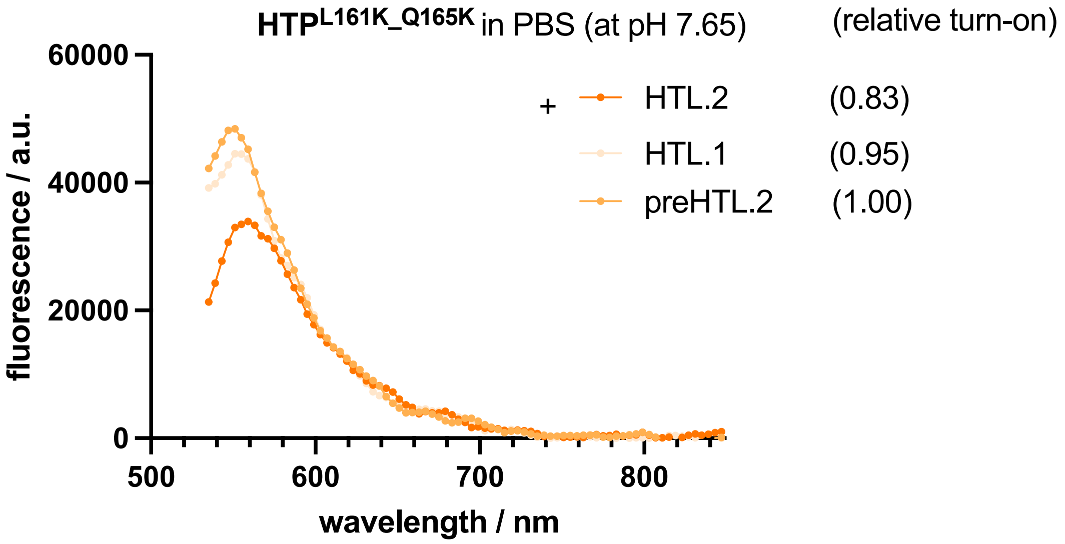

**Supplementary Fig. 4: Influence of HTL on the binary system.** Raw fluorescence of 2.5 µM of $F_2$X-HTL.1, $F_2$X-HTL.2 or $F_2$X-preHTL.2 and HTP$^{L161K_Q165K}$ mutant in PBS after 16 h incubation at 37 °C at pH = 7.65 (n = 3). AUC [Total Area (normalized): HTL.2: 2631732 (83%), HTL.1: 3030795 (95%), preHTL.2: 3189648 (100%)]

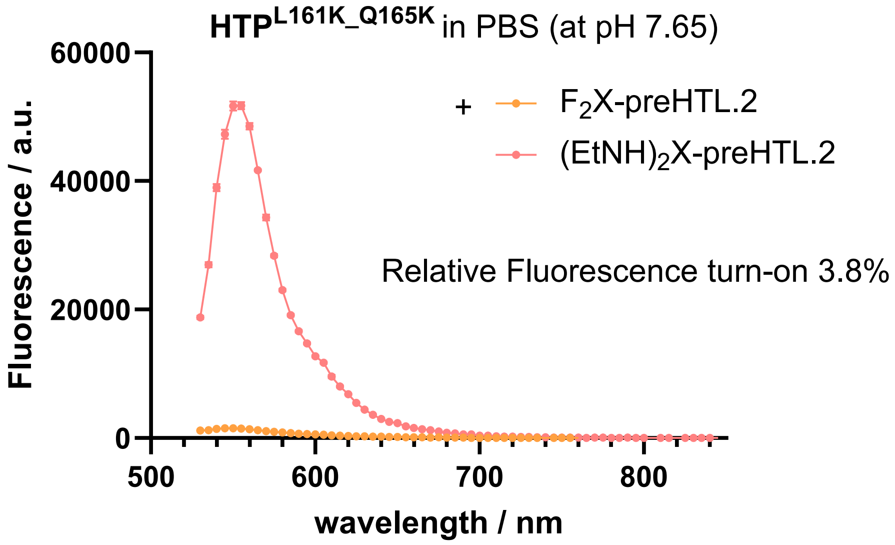

**Supplementary Fig. 5: Relative efficiency of system.** Raw fluorescence of 2.5 µM of $F_2X$-preHTL.2 or $(EtNH)_2X$-preHTL.2 and HTP$^{L161K_Q165K}$ mutant in PBS after 16 h incubation at 37 °C, at pH = 7.65. (n = 3). Relative fluorescent turn-on was calculated by AUC ($F_2X$) / AUC ($(EtNH_2)X$).

### Supplementary Scheme

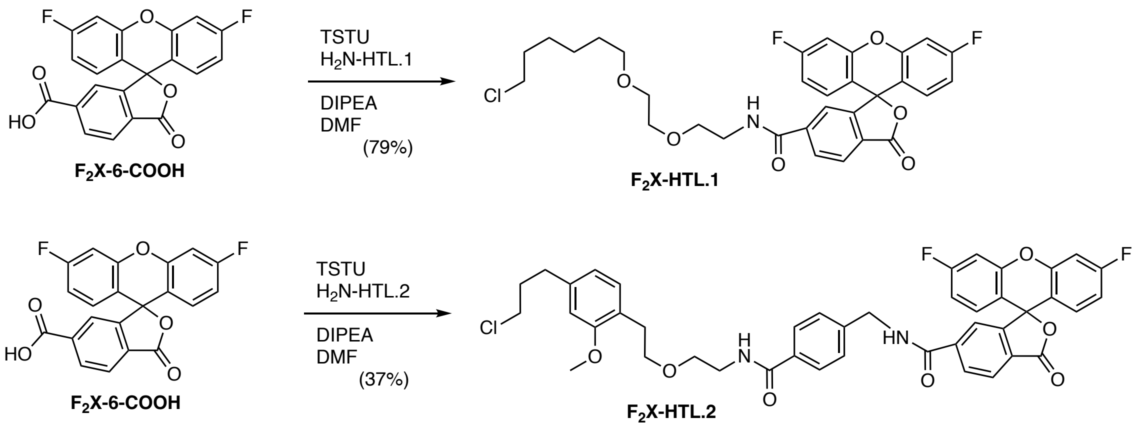

**Supplementary Scheme 1: Synthesis of F$_2$X-HTL.1 and F$_2$X-HTL.2.**

### Methods

#### General UV/Vis and fluorescence spectroscopy

All dyes for spectroscopy were prepared as stock solutions in DMSO-d6, concentration was determined by $^{1}$H qNMR in presence of DMF as internal standard. These solutions were diluted such that the final DMSO concentration did not exceed 1% v/v. All measurements were taken at ambient temperature (22 ± 2 °C). Absorption spectra were recorded on a Cary Model 100 spectrometer (Agilent), and fluorescence spectra were recorded on a Cary Eclipse fluorometer (Varian). The spectra, maximum absorption wavelength ($\lambda_{abs}$), and maximum emission wavelength ($\lambda_{em}$) were measured in PBS, pH 7.4 buffer or EtOH + 1% TFA (n = 3). Normalized spectra are shown for clarity. Turn-on was measured on a M200Pro plate reader (TECAN) with $\lambda_{exc}$ = 500 nm at 25 °C. Concentrations were [$F_2X$] = 2.5 uM, [$(EtNH)_2X$] = 200 nM, [HTP mutants] = 400 nM.

#### Quantum Yield Determination

Absolute fluorescence quantum yields (Φ) were measured using Quantaurus-QY Absolute PL (HAMAMATSU, model C11347-11) with a 150 W xenon light source using a clear window quartz cuvette with long quartz tube mouth (10 mm light path) from Hamamatsu. This instrument uses an integrating sphere to determine photons absorbed and emitted by a sample. Measurements were carried out using dilute samples (A < 0.1) and self-absorption corrections were performed using the instrument software. We validated the quantum yield values using fluorophores with established quantum yield values (i.e. fluorescein in 0.1 M NaOH; measured: 0.88; literature: 0.86– 0.92).

#### Statistical analysis and graphical design

All statistical tests were performed using GraphPad Prism version 9.3, 10 or 11 software and are listed in the figure legends. Normal distribution of data sets was tested with one-way ANOVA and Tukey's multiple comparisons test or two-way ANOVA and Šídák's multiple comparisons test. For all experiments statistical significance was assigned, with an alpha-level of $p < 0.05$. Figures were assembled using MS Office (version 16) and PyMol 3.

#### Cloning

The Halo tag Protein was mutated using the Q5 Site-Directed Mutagenesis Kit (NEB, E0554S). The primers were designed using the NEB Base Changer tool and ordered from Sigma Aldrich. Positive PCR products were transformed into chemically competent bacteria *E. coli* DH5$\alpha$ (NEB, C2987H) following KLD reaction. Plasmids were sequenced at Eurofins using appropriate primers. The resulting sequences were aligned against the original sequence using Emboss needle pairwise sequence alignment tool.

#### Protein expression and purification

HTP was expressed and purified as described previously (ref[1]): HTP with an N-terminal Strep-tag and C-terminal 10xHis-tag was cloned into a pET51b(+) expression vector for bacterial expression. For purification, HTP mutants were expressed in *E. coli* strain BL21 DE3. LB media contained ampicillin (100 mg $mL^{-1}$, Carl Roth, K029.2) for protein expression. A culture was grown at 37 °C until an $OD_{600}$ of 0.6 was reached at which point cells were induced with IPTG (1 mM, Carl Roth, CN08.1). Protein constructs were expressed overnight at 16 C. Cells were harvested by centrifugation and sonicated to produce cell lysates. The lysate was cleared by centrifugation and purified by NiNTA resin (Thermo Fisher, 88222) according to the manufacturer's protocols. Purified protein samples were aliquoted in PBS, flash frozen and stored at –80 C.

#### Full protein mass spectrometry

Labelling substrates were dissolved in DMSO to a concentration of 1 mM and diluted in activity buffer (containing: 50 mM NaCl, 50 mM HEPES, pH=7.3) to 20 µM. Protein was diluted in activity buffer to a concentration of 2 µM. 25 µL of each protein and labelling agent were combined and allowed to incubate at room temperature for the indicated time period, before full protein mass was acquired. Full protein MS was measured using a Waters XEVO G2-XS quadrupole time-of-flight (Q-TOF) mass spectrometer (Waters cooperation, USA) alongside an, Acquity UPLC system with an Acquity UPLC Protein BEH C4 column (1.7 µm, 2.1 mm x 50 mm). Samples were eluted with a flow rate of 0.4 mL/min. The following gradient was used: A: 0.01% FA in $H_2O$; B: 0.01 % FA in MeCN. 5-85% B: 0–3.5 min; 85-5% B: 3.7-4 min; 5-85% B: 4-4.5 min; 85-5% B: 4.5-5.5 min. MS was acquired over the range of 500-4000 Da using continuous scanning with 1 s scan time and a cone voltage of 80 V with subsequent spectral deconvolution using the MaxEnt1 algorythm at a resolution of 1 Da/channel until convergence. Samples were measured n = 1. All samples were prepared as 0.1 mg/mL solutions in PBS.

#### Fluorescence polarization

Fluorescence polarization measurements were performed on a TECAN Spark Cyto and on a TECAN GENios Pro plate reader. Samples from full protein mass spectrometry experiments were used and transferred into a Greiner black flat bottom 96 well plate. $\lambda_{Ex}$ = 535±25 nm; $\lambda_{Em}$ = 590±35 nm;10 flashes; 40ms integration time.

#### Cell culture

HEK293T cells were cultured in growth media (DMEM, Glutamax, 4.5 g Glucose, 10% FCS, 1% PS; Invitrogen, 10566016) at 37 °C and 5% $CO_2$. 50,000 cells per well were seeded on 8-well *µ*L slides (Ibidi, 80826) previously coated with poly-L-lysine (Aldrich, mol wt 70 000–150 000, P6286). The next day, 400 ng DNA was transfected using 0.8 *µ*L Jet Prime reagent in 40 *µ*L Jet Prime buffer (VWR, 101000046) per well/plasmid. Media was exchanged against antibiotic-free media before the transfection mix was applied on the cells. After 4 hours incubation at 37 °C and 5% $CO_2$, medium was exchanged against growth media containing 2.5 *µ*M $F_2$X-preHTL.2 or $F_2$X-preSHTL. Cells were incubated over night and stained next day with 1 *µ*M BG-SiR-d12 and 1 *µ*M Hoechst for 30 min at 37°C. Cells were washed once in fluorobrite (Invitrogen, A1896701) and imaged in fluorobrite.

#### Confocal microscopy

For confocal imaging cells were imaged using the NIKON-CSU-X1 (Nikon Ti Eclipse with automatic stage and Perfect Focus System, equipped with Yokogawa spinning disk (CSU-X1, 1000 scan/s) and EMCCD Camera (AU-888, 13 um pixel) and laser for Hoechst $\lambda$ = 405 nm, $F_2$X-preHTL.2 or $F_2$X-preSHTL $\lambda$ = 514 nm and BG-SiR-d12 / BG-Sulfo646 $\lambda$ = 638 nm for fast confocal imaging, objectives: 40x air NA 0.95 in a temperature-controlled chamber at 37° C. For the bleaching experiment, cells were used as prepared for confocal microscopy. To induce bleaching 100 % laser power $\lambda$ = 514 nm was used for $F_2$X-preHTL.2 and Et$(NH)_2$-preHTL. The resulting images were analyzed using Fiji software. Integrated density was measured for 514 nm and 638 nm, normalized to t=0 and plotted in GraphPad Prism 10.

#### Single molecule spectroscopy

Samples for no-wash protein labelling were prepared as follows:

stocks:

| | |
|---|---|
| $F_2$X-preHTL.2 | 2.4 mM stock in DMSO |
| $(EtNH)_2$X-preHTL.2 | 2.0 mM stock in DMSO |
| $HTP^{L161K\_Q165K}$ | 579 µM |
| $HTP^{L161A\_Q165A}$ | 395 µM |

HTP was labelled in PBS (pH = 8.65) o.n. at r.t. at the following concentrations with [HTP] = 5 µM and [dye-preHTL.2] = 10 µM in PCR tubes with a total volume of 20 µL. The next day, samples were diluted 1:10,000 in PBS pH = 7.4 into 8 well ibidi dished.

Fluorescence lifetimes and fluorescence correlation spectroscopy were recorded on a Luminosa confocal single molecule sensitive spectrometer (Picoquant, Berlin). Fluorophores were excited using a linearly polarized pulsed laser with 485 nm wavelength (LDH-D-C-485S, Picoquant), operated through a Sepia PDL 828-L laser driver (Picoquant) at a pulse rate of 20 MHz and through a 60x water immersion objective (UPLSAPO 60 x ultra-planapochromat, Olympus). The fluorescence light was then collected through a pinhole of 20 µm diameter, filtered with an HC 520/35 emission filter (Chroma) and split into parallel and perpendicular detection channels. Photons were detected on two Single Photon Counting Modules (SPADs, Hamamatsu) and counted using a MultiHarp 150 4P (Picoquant). Fluorescence anisotropy was calculated using the following equation:

For recording fluorescence lifetimes and fluorescence correlation curves, no polarizing beam splitter was used in the emission path. Fluorescence lifetimes decays and fluorescence correlation curves were calculated and fit using the software SymPhoTime (Picoquant).

### Protein sequences

```
HTP               MASWSHPQFEKGADDDDKVPHGSEIGTGFPFDPHYVEVLGERMHYVDVGPRDGTPVLFLH     60
HTP_L161A_Q165A   MASWSHPQFEKGADDDDKVPHGSEIGTGFPFDPHYVEVLGERMHYVDVGPRDGTPVLFLH     60
HTP_L161K_Q165K   MASWSHPQFEKGADDDDKVPHGSEIGTGFPFDPHYVEVLGERMHYVDVGPRDGTPVLFLH     60
HTP_Q165K_E170K   MASWSHPQFEKGADDDDKVPHGSEIGTGFPFDPHYVEVLGERMHYVDVGPRDGTPVLFLH     60
HTP_E170K_M175K   MASWSHPQFEKGADDDDKVPHGSEIGTGFPFDPHYVEVLGERMHYVDVGPRDGTPVLFLH     60
HTP_A151K_E174K   MASWSHPQFEKGADDDDKVPHGSEIGTGFPFDPHYVEVLGERMHYVDVGPRDGTPVLFLH     60
HTP_A151K_L161K   MASWSHPQFEKGADDDDKVPHGSEIGTGFPFDPHYVEVLGERMHYVDVGPRDGTPVLFLH     60
HTP_F144K_M175K   MASWSHPQFEKGADDDDKVPHGSEIGTGFPFDPHYVEVLGERMHYVDVGPRDGTPVLFLH     60
HTP_T148K_V167K   MASWSHPQFEKGADDDDKVPHGSEIGTGFPFDPHYVEVLGERMHYVDVGPRDGTPVLFLH     60

HTP               GNPTSSYVWRNIIPHVAPTHRCIAPDLIGMGKSDKPDLGYFFDDHVRFMDAFIEALGLEE     120
HTP_L161A_Q165A   GNPTSSYVWRNIIPHVAPTHRCIAPDLIGMGKSDKPDLGYFFDDHVRFMDAFIEALGLEE     120
HTP_L161K_Q165K   GNPTSSYVWRNIIPHVAPTHRCIAPDLIGMGKSDKPDLGYFFDDHVRFMDAFIEALGLEE     120
HTP_Q165K_E170K   GNPTSSYVWRNIIPHVAPTHRCIAPDLIGMGKSDKPDLGYFFDDHVRFMDAFIEALGLEE     120
HTP_E170K_M175K   GNPTSSYVWRNIIPHVAPTHRCIAPDLIGMGKSDKPDLGYFFDDHVRFMDAFIEALGLEE     120
HTP_A151K_E174K   GNPTSSYVWRNIIPHVAPTHRCIAPDLIGMGKSDKPDLGYFFDDHVRFMDAFIEALGLEE     120
HTP_A151K_L161K   GNPTSSYVWRNIIPHVAPTHRCIAPDLIGMGKSDKPDLGYFFDDHVRFMDAFIEALGLEE     120
HTP_F144K_M175K   GNPTSSYVWRNIIPHVAPTHRCIAPDLIGMGKSDKPDLGYFFDDHVRFMDAFIEALGLEE     120
HTP_T148K_V167K   GNPTSSYVWRNIIPHVAPTHRCIAPDLIGMGKSDKPDLGYFFDDHVRFMDAFIEALGLEE     120

HTP               VVLVIHDWGSALGFHWAKRNPERVKGIAFMEFIRPIPTWDEWPEFARETFQAFRTTDVGR     180
HTP_L161A_Q165A   VVLVIHDWGSALGFHWAKRNPERVKGIAFMEFIRPIPTWDEWPEFARETFQAFRTTDVGR     180
HTP_L161K_Q165K   VVLVIHDWGSALGFHWAKRNPERVKGIAFMEFIRPIPTWDEWPEFARETFQAFRTTDVGR     180
HTP_Q165K_E170K   VVLVIHDWGSALGFHWAKRNPERVKGIAFMEFIRPIPTWDEWPEFARETFQAFRTTDVGR     180
HTP_E170K_M175K   VVLVIHDWGSALGFHWAKRNPERVKGIAFMEFIRPIPTWDEWPEFARETFQAFRTTDVGR     180
HTP_A151K_E174K   VVLVIHDWGSALGFHWAKRNPERVKGIAFMEFIRPIPTWDEWPEFARETFQKFRTTDVGR     180
HTP_A151K_L161K   VVLVIHDWGSALGFHWAKRNPERVKGIAFMEFIRPIPTWDEWPEFARETFQKFRTTDVGR     180
HTP_F144K_M175K   VVLVIHDWGSALGFHWAKRNPERVKGIAFMEFIRPIPTWDEWPEKARETFQAFRTTDVGR     180
HTP_T148K_V167K   VVLVIHDWGSALGFHWAKRNPERVKGIAFMEFIRPIPTWDEWPEFAREKFQAFRTTDVGR     180
                  ******************************************** *** ** ********

HTP               KLIIDQNVFIEGTLPMGVVRPLTEVEMDHYREPFLNPVDREPLWRFPNELPIAGEPANIV     240
HTP_L161A_Q165A   AIIDANVFIEGTLPMGVVRPLTEVEMDHYREPFLNPVDREPLWRFPNELPIAGEPANIV      240
HTP_L161K_Q165K   KKIIDKNVFIEGTLPMGVVRPLTEVEMDHYREPFLNPVDREPLWRFPNELPIAGEPANIV     240
HTP_Q165K_E170K   KLIIDKNVFIKGTLPMGVVRPLTEVEMDHYREPFLNPVDREPLWRFPNELPIAGEPANIV     240
HTP_E170K_M175K   KLIIDQNVFIKGTLPKGVVRPLTEVEMDHYREPFLNPVDREPLWRFPNELPIAGEPANIV     240
HTP_A151K_E174K   KLIIDQNVFIEGTLPMGVVRPLTEVEMDHYREPFLNPVDREPLWRFPNELPIAGEPANIV     240
HTP_A151K_L161K   KLIIDKNVFIEGTLPMGVVRPLTEVEMDHYREPFLNPVDREPLWRFPNELPIAGEPANIV     240
HTP_F144K_M175K   KLIIDQNVFIEGTLPKGVVRPLTEVEMDHYREPFLNPVDREPLWRFPNELPIAGEPANIV     240
HTP_T148K_V167K   KLIIDQNKFIEGTLPMGVVRPLTEVEMDHYREPFLNPVDREPLWRFPNELPIAGEPANIV     240
                    *** * ** **** ********************************************

HTP               ALVEEYMDWLHQSPVPKLLFWGTPGVLIPPAEAARLAKSLPNCKAVDIGPGLHLLQEDNP     300
HTP_L161A_Q165A   ALVEEYMDWLHQSPVPKLLFWGTPGVLIPPAEAARLAKSLPNCKAVDIGPGLHLLQEDNP     300
HTP_L161K_Q165K   ALVEEYMDWLHQSPVPKLLFWGTPGVLIPPAEAARLAKSLPNCKAVDIGPGLHLLQEDNP     300
HTP_Q165K_E170K   ALVEEYMDWLHQSPVPKLLFWGTPGVLIPPAEAARLAKSLPNCKAVDIGPGLHLLQEDNP     300
HTP_E170K_M175K   ALVEEYMDWLHQSPVPKLLFWGTPGVLIPPAEAARLAKSLPNCKAVDIGPGLHLLQEDNP     300
HTP_A151K_E174K   ALVEEYMDWLHQSPVPKLLFWGTPGVLIPPAEAARLAKSLPNCKAVDIGPGLHLLQEDNP     300
HTP_A151K_L161K   ALVEEYMDWLHQSPVPKLLFWGTPGVLIPPAEAARLAKSLPNCKAVDIGPGLHLLQEDNP     300
HTP_F144K_M175K   ALVEEYMDWLHQSPVPKLLFWGTPGVLIPPAEAARLAKSLPNCKAVDIGPGLHLLQEDNP     300
HTP_T148K_V167K   ALVEEYMDWLHQSPVPKLLFWGTPGVLIPPAEAARLAKSLPNCKAVDIGPGLHLLQEDNP     300

HTP               DLIGSEIARWLSTLEISGAPGFSSISAHHHHHHHHHH          337
HTP_L161A_Q165A   DLIGSEIARWLSTLEISGAPGFSSISAHHHHHHHHHH          337
HTP_L161K_Q165K   DLIGSEIARWLSTLEISGAPGFSSISAHHHHHHHHHH          337
HTP_Q165K_E170K   DLIGSEIARWLSTLEISGAPGFSSISAHHHHHHHHHH          337
HTP_E170K_M175K   DLIGSEIARWLSTLEISGAPGFSSISAHHHHHHHHHH          337
HTP_A151K_E174K   DLIGSEIARWLSTLEISGAPGFSSISAHHHHHHHHHH          337
HTP_A151K_L161K   DLIGSEIARWLSTLEISGAPGFSSISAHHHHHHHHHH          337
HTP_F144K_M175K   DLIGSEIARWLSTLEISGAPGFSSISAHHHHHHHHHH          337
HTP_T148K_V167K   DLIGSEIARWLSTLEISGAPGFSSISAHHHHHHHHHH          337
```

### Molecular Modelling

#### Structure Preparation and Covalent Docking

The initial geometry of the receptor protein was obtained from the Protein Data Bank (entry PDB-5UY1 for apoHTP).[2] The model was protonated for physiological conditions using OpenBabel 3.1 and used directly for subsequent docking.[3]

The ligand structure was generated from SMILES representation using OpenBabel 3.1 and converted to three-dimensional geometry with standard protonation at physiological conditions and using the conformer generation setting “Best”. The structure was saved in SDF format and used without further modification.

Covalent docking was performed using GNINA 1.3.[4–6] The full protein structure was used as the receptor, and the docking search space was defined automatically using the entire receptor with additional padding of 5.0 Å and automatic box extension enabled. Covalent attachment was specified between the carboxylate of residue Asp106 and the respective terminal carbon atom as a single bond. Flexibility of the receptor was introduced locally by allowing side-chain flexibility for the covalent aspartate.

Docking was performed using the Vinardo scoring function, followed by convolutional neural network (CNN)-based rescoring with the general_default2018 parametrization. The search exhaustiveness was set to 128, and the random seed was set to 100. The docked poses were ranked by the CNN confidence score, and the highest scoring complex was used as starting point for subsequent molecular dynamics simulations.

#### Molecular Dynamics Simulation

All molecular dynamics simulations were performed using GROMACS 2025.4.[7–14] First, the protein structure was protonated for physiological conditions using PDBFixer, a tool supplied within the OpenMM 8.2 suite.[15] Subsequently, the protein was parametrized using the AMBER14SB protein force field including improper dihedrals, as supplied in the OpenFF toolbox.[16–18] All non-standard bonds and atoms were parametrized using the OpenFF Sage 2.3 force field.[16] The system was solvated in a dodecahedral box of TIP3P water with a minimum solute-box distance of 10 Å and neutralized with NaCl to a final concentration of 150 mM.

Periodic boundary conditions were applied in all dimensions. Short-range electrostatic and van-der-Waals interactions were truncated at 1.0 nm, and long-range electrostatics were treated using the particle mesh Ewald (PME) method.[19] Neighbour searching was performed using the Verlet cutoff scheme with an update frequency of 20 steps. Analytical dispersion corrections to energy and pressure were applied.

Energy minimization was carried out using the steepest descent algorithm for up to 50,000 steps with an initial step size of 0.01 nm and a convergence cutoff of 500.0 kJ $mol^{-1}$ $nm^{-1}$. The minimized system was first equilibrated in the NVT ensemble for 5 ns at 300 K using a 1 fs integration time step. Temperature coupling was achieved using the velocity-rescaling thermostat (V-rescale) with a coupling constant of 0.1 ps. Subsequently, the systems were equilibrated in the NPT ensemble for 5 ns at 300 K and 1 bar using the Parinello-Rahman barostat (coupling constant of 2.0 ps, compressibility 4.5e-5 $bar^{-1}$).[20]

All bonds involving hydrogen atoms were constrained using the LINCS algorithm (order 4, one iteration).[21] Production simulations were then performed in the NPT ensemble for 100 ns using a 2.0 fs integration time step under identical thermostat and barostat settings. Coordinates were written every 100 ps during production, unwrapping and alignment of the trajectories was performed using GROMACS.

#### Target Residue Sampling

For each frame of the aligned and unwrapped MD production trajectory, protein residues were reduced to the Cartesian centroid of their heavy side-chain atoms using a custom Python script. Backbone atoms and hydrogen atoms were excluded from the centroid definition. Glycine residues were not considered unless stated otherwise, as they do not contain a side chain. For each residue $i$ and trajectory frame $t$, the minimum distance $d_i(t)$ between the side-chain centroid and either of the two fluoride-bearing carbon atoms of the covalently linked $F_2X$ ligand was calculated.

Residues were evaluated using two complementary criteria. First, a cutoff-based contact occupancy was calculated as the fraction of trajectory frames in which $d_i(t)$ was below the chosen proximity threshold of $d_{\text{ref}} = 8\,\text{Å}$. Residues exceeding 10% contact occupancy were considered proximity-enriched candidate positions. Second, to obtain a cutoff-free ranking that accounts for persistent close approaches throughout the full trajectory, an inverse-distance score $S_i$ was calculated for every residue. This score weights shorter centroid-electrophile distances more strongly while still retaining contributions from all frames.

The inverse-distance score was defined as:

$$S_i = \frac{1}{T}\sum_{t=1}^{T}\left[\frac{d_{\text{ref}}}{\min\limits_{j\in\{1,2\}} \left\| \left(\frac{1}{N_i}\sum_{a=1}^{N_i} r_{i,a}(t)\right) - x_j(t) \right\|_2}\right]^2, \qquad (d_{\text{ref}} = 8\,\text{Å})$$

Here, $\boldsymbol{T}$ is the total number of analyzed trajectory frames, $\boldsymbol{N_i}$ is the number of heavy side-chain atoms of residue $\boldsymbol{i}$, $\boldsymbol{r_{i,a}(t)}$ is the Cartesian coordinate of side-chain atom $\boldsymbol{a}$ of residue $\boldsymbol{i}$ in frame $\boldsymbol{t}$, and $\boldsymbol{x_j(t)}$ is the Cartesian coordinate of fluoride-bearing carbon atom $\boldsymbol{j}$ of the $F_2X$ ligand. For each residue and frame, the shorter of the two distances to the electrophilic xanthene carbon atoms was used.

The cutoff-based occupancy and the inverse-distance score were used jointly to prioritize residues that repeatedly sampled spatial proximity to the reactive xanthene positions. Candidate sites were subsequently inspected structurally, with emphasis on surface accessibility, compatibility with lysine substitution, and potential for pairwise positioning around the two electrophilic carbon atoms.

### Synthesis

#### General chemistry

Chemicals and solvents were purchased from Merck (Merck group, Germany), TCI (Tokyo chemical industry CO., LTD., Japan) and Acros Organics (Thermo Fisher scientific, USA) and used without further purification. Dry solvents were purchased from Acros Organics (Thermo Fisher scientific, USA).

UPLC-UV/Vis for purity assessment was performed on an Agilent 1260 Infinity II LC System equipped with Agilent SB-C18 column (1.8 µm, 2.1 × 50 mm). Buffer A: 0.1% FA in $H_2O$ Buffer B: 0.1% FA acetonitrile. The typical gradients were:
-from 10% B for 1.0 min→ gradient to 95% B over 5 min→ 95% B for 1.0 min with (Method A).
-from 30% B for 1.0 min→ gradient to 95% B over 5 min→ 95% B for 1.0 min with (Method B).
-from 50% B for 1.0 min→ gradient to 95% B over 5 min→ 95% B for 1.0 min with (Method C).
Retention times ($t_R$) are given in minutes (min). Chromatograms were imported into GraphPad Prism 8 and purity was determined by calculating AUC ratios.

Preparative or semi-preparative HPLC was performed on an Agilent 1260 Infinity II LC System equipped with columns as followed: preparative column – Reprospher 100 C18 columns (10 µm: 50 x 30 mm at 20 mL/min flow rate; semi-preparative column – 5 µm: 250 x 10 mm at 4 mL/min flow rate. Eluents A (0.1% TFA in $H_2O$) and B (0.1% TFA in MeCN) were applied as a linear gradient. Peak detection was performed at maximal absorbance wavelength.

For HRMS, samples were analyzed on Orbitrap Fusion mass spectrometer (Thermo Fisher Scientific). MS scans were acquired in a range of 350 to 1500 m/z. MS1 scans were acquired in the Orbitrap with a mass resolution of 120,000 with an AGC target value of 4e5 and 50 ms injection time. MS2 scans were acquired in the ion trap with an AGC target value of 1e4 and 35 ms injection time. Precursor ions with charge states 2-4 were isolated with an isolation window of 1.6 m/z and 40 sec dynamic exclusion. Precursor ions were fragmented using higher-energy collisional dissociation (HCD) with 30% normalized collision energy.

#### 3',6'-Difluoro-3-oxo-3*H*-spiro[isobenzofuran-1,9'-xanthene]-6-carboxylic acid, $F_2$X-6-COOH

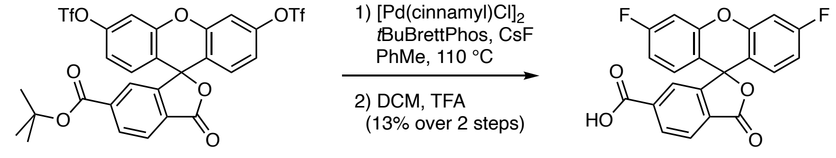

A Schlenk tube was charged with cesium fluoride (131 mg, 0.861 mmol, 10 equiv.). The salt was dried under high vacuum at 150 °C during 2h prior to use under $N_2$ and store in a sealed tube. After cooling to r.t., 6-*tert*-butoxycarbonylfluorescein ditriflate **1** (60.0 mg, 86.1 µmol), $[Pd(cinnamyl)Cl]_2$ (4.5 mg, 0.861 µmol, 0.1 equiv.), *t*BuBrettPhos (6.3 mg, 12.9 µmol, 0.15 equiv.) were added and dissolved with *dry* toluene under $N_2$. The Schlenk tube was sealed and evacuated/backfilled with $N_2$ (3×). The reaction mixture was stirred at 110 °C during 14h. After cooling to r.t., the crude mixture was filtrated through a plug of Celite and washed with MeOH than concentrated under vacuum. The crude residue was then diluted in DCM/TFA (5.0 mL, 3:1) and stirred during 4 h. The crude material was concentrated to dryness. The residue obtained was dissolved in a DMSO:$H_2O$:MeCN:AcOH (10:25:25:1) and subjected to RP-HPLC (MeCN:$H_2O$+0.1% TFA = 30:70 to 95:05 over 40 minutes), to afford 4.2 mg (13%, TFA salt) of **1** as a pale yellow solid.

**$^1$H NMR** (600 MHz, MeOD-d4) δ 8.35 (dd, *J* = 8.0, 1.2 Hz, 1H), 8.14 (d, *J* = 8.0 Hz, 1H), 7.77 (s, 1H), 7.19 (d, *J* = 9.5, 2.3 Hz, 2H), 6.97 – 6.89 (m, 4H).
**$^{13}$C NMR** (151 MHz, MeOD-d4) δ 169.7 (C), 167.7 (C), 165.2 (C, d, *J* = 249 Hz), 154.2 (C), 153.4 (C, d, *J* = 12.8 Hz), 139.3 (C), 132.8 (CH), 131.2 (CH, d, *J* = 10.4 Hz), 130.8 (C), 126.5 (CH), 126.1 (CH), 116.2 (C, d, *J* = 2.9 Hz), 113.4 (CH, d, *J* = 22.8 Hz), 105.2 (CH, d, *J* = 25.9 Hz), 83.2 (C).
**$^{19}$F NMR** (564 MHz, MeOD-d4) δ -77.0, -110.4 ($m_C$, *J* = 7.8 Hz).
**HRMS** (ESI): calc. for $C_{21}H_{11}F_2O_5$ $[M+H]^+$: 381.0570, found: 381.0532.

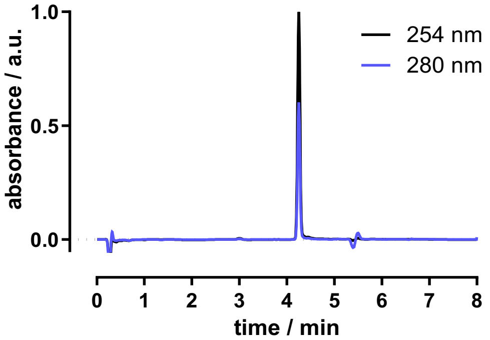

#### *N*-(2-(4-(3-Chloropropyl)-2-methoxyphenethoxy)ethyl)-3',6'-difluoro-3-oxo-3*H*-spiro[isobenzofuran-1,9'-xanthene]-6-carboxamide, F₂X-HTL.1

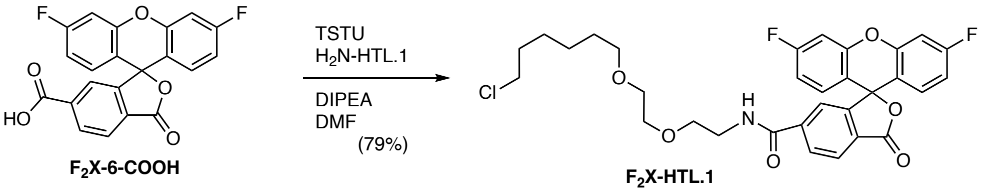

An Eppendorf tube was charged with **$F_2$X-6-COOH** (1.00 mg, 2.63 μmol, 1.0 equiv.) in DMF (100 μL). DIPEA (3.6 μL, 21.0 μmol, 8.0 equiv.) and a 100 mM solution of TSTU (39.4 μL, 3.94 μmol, 1.5 equiv.) in DMF were added successively. The reaction mixture was allowed to incubate for 15 minutes, and then HTL.1-$NH_2$ (1.0 mg, 3.84 μmol, 1.5 equiv.) was added in one drop. The reaction mixture was incubated at r.t. until complete conversion was observed (by LCMS). The mixture was diluted with $H_2O$:MeCN:AcOH (25:25:1) and subjected to RP-HPLC (MeCN:$H_2O$+0.1% TFA = 50:90 to 95:05 over 40 minutes) to obtain 1.45 mg (79%, TFA salt) of **$F_2$X-HTL.1** after lyophilization as a white solid.

**$^1$H NMR** (600 MHz, MeOD-d4) δ 8.18 (dd, *J* = 8.1, 1.3 Hz, 1H), 8.13 (d, *J* = 8.0 Hz, 1H), 7.66 (s, 1H), 7.21 – 7.16 (m, 2H), 6.93 (dd, *J* = 7.6, 1.6 Hz, 4H), 3.61 – 3.55 (m, 4H), 3.55 – 3.48 (m, 5H), 3.40 (t, *J* = 6.5 Hz, 2H), 1.76 – 1.68 (m, 2H), 1.49 (q, *J* = 6.7 Hz, 2H), 1.43 – 1.39 (m, 2H), 1.36 – 1.31 (m, 2H).

**$^{19}$F NMR** (564 MHz, MeOD-d4) δ -77.3, -110.5 ($m_C$, *J* = 7.4 Hz).

**HRMS** (ESI): calc. for $C_{35}H_{31}ClF_2NO_6$ $[M+H]^+$: 586.1803, found: 586.1815.

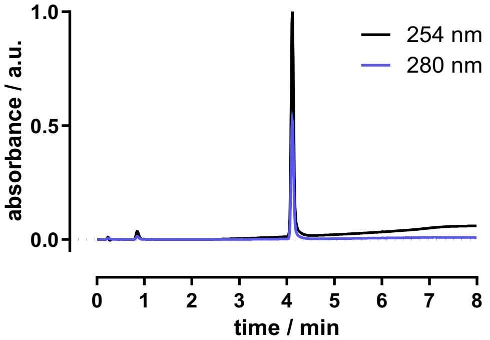

#### *N*-(2-(4-(3-Chloropropyl)-2-methoxyphenethoxy)ethyl)-3',6'-difluoro-3-oxo-3*H*-spiro[isobenzofuran-1,9'-xanthene]-6-carboxamide, $F_2$X-preHTL.2

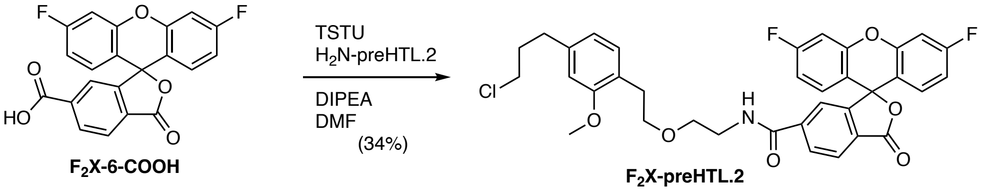

An Eppendorf tube was charged with **$F_2$X-6-COOH** (1.50 mg, 3.94 µmol, 1.0 equiv.) in DMF (100µL). DIPEA (5.5 µL, 31.6 µmol, 8.0 equiv.) and a 100 mM solution of TSTU (59 µL, 5.92 µmol, 1.5 equiv.) in DMF were added successively. The reaction mixture was allowed to incubate for 15 minutes, and then a 100 mM preHTL.2-$NH_2$ solution (59.2 µL, 5.92 µmol, 1.5 equiv.) was added in one drop. The reaction mixture was incubated at r.t. until complete conversion was observed (by LCMS). The mixture was diluted with $H_2O$:MeCN:AcOH (25:25:1) and subjected to RP-HPLC (MeCN:$H_2O$+0.1% TFA = 50:90 to 95:05 over 40 minutes) to obtain 1.00 mg (34%, TFA salt) of **$F_2$X-preHTL.2** after lyophilization as a white solid.

**$^1$H NMR** (600 MHz, MeOD-d4) δ 8.1 (s, 2H), 7.6 (s, 1H), 7.2 (d, *J* = 9.5 Hz, 2H), 7.0 (d, *J* = 7.5 Hz, 1H), 6.9 (dd, *J* = 7.0, 1.3 Hz, 4H), 6.7 (s, 1H), 6.6 (d, *J* = 7.5 Hz, 1H), 3.8 (s, 3H), 3.6 (t, *J* = 7.1 Hz, 2H), 3.5 (t, *J* = 5.6 Hz, 2H), 3.5 (t, *J* = 6.5 Hz, 2H), 3.5 (t, *J* = 5.5 Hz, 2H), 2.8 (t, *J* = 7.1 Hz, 2H), 2.7 – 2.7 (m, 2H), 2.0 (dt, *J* = 13.6, 6.6 Hz, 2H).

**$^{19}$F NMR** (564 MHz, MeOD-d4) δ -77.3, -110.5 ($m_C$, *J* = 7.4 Hz).

**$^{13}$C NMR** (151 MHz, MeOD-d4) δ 169.8 (C), 168.1 (C), 164.3 (C, d, *J* = 249 Hz), 159.1 (C), 154.5 (C), 153.4 (C, d, *J* = 12.2 Hz), 142.9 (C), 141.9 (C), 131.5 (CH), 131.4 (CH, d, *J* = 9.9 Hz), 130.8 (CH), 129.5 (C), 126.5 (CH), 125.7 (C), 123.9 (CH), 121.4 (CH), 116.2 (C, d, *J* = 1.9 Hz),113.5 (CH, d, *J* = 22.9 Hz), 111.8 (CH), 105.28 (CH, d, *J* = 25.8 Hz), 71.7 ($CH_2$), 69.7 ($CH_2$), 55.7 ($CH_3$-O), 45.0 ($CH_2$), 41.1 ($CH_2$), 35.4 ($CH_2$), 33.8 ($CH_2$), 31.3 ($CH_2$). (One Cq is missing).

**HRMS** (ESI): calc. for $C_{35}H_{31}ClF_2NO_6$ $[M+H]^+$: 634.1803, found: 634.1788.

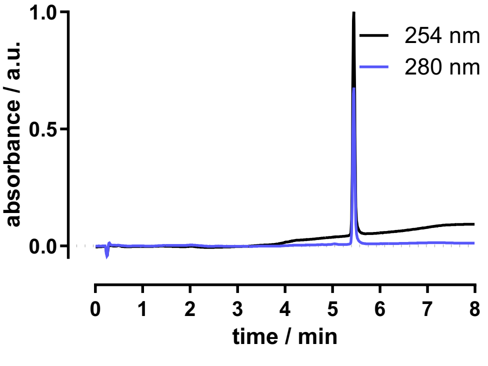

#### *N*-(4-((2-(4-(3-Chloropropyl)-2-methoxyphenethoxy)ethyl) carbamoyl)benzyl)-3',6'-difluoro-3-oxo-3H-spiro[isobenzofuran-1,9'-xanthene]-6-carboxamide, $F_2$X-HTL.2

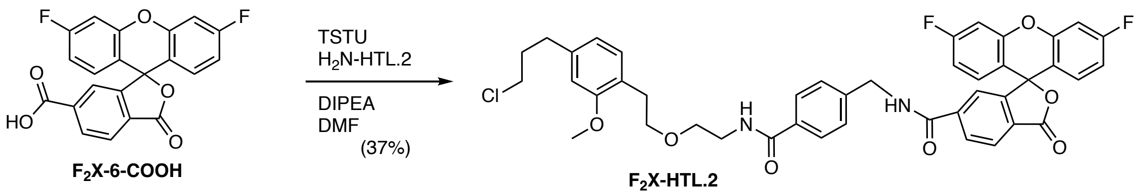

An Eppendorf tube was charged with **$F_2$X-6-COOH** (0.70 mg, 1.80 µmol, 1.0 equiv.) in DMF (100µL). DIPEA (2.5 µL, 15.0 µmol, 8.0 equiv.) and a 100 mM solution of TSTU (37 µL, 3.70 µmol, 2.0 equiv.) in DMF were added successively. The reaction mixture was allowed to incubate for 15 minutes, and then a 100m M HTL.2-$NH_2$ solution (37 µL, 3.70 µmol, 2.0 equiv.) was added in one drop. The reaction mixture was incubated at r.t. until complete conversion was observed (by LCMS). The mixture was diluted with $H_2O$:MeCN:AcOH (25:25:1) and subjected to RP-HPLC (MeCN:$H_2O$+0.1% TFA = 50:90 to 95:05 over 40 minutes) to obtain 0.60 mg (37%, TFA salt) of **$F_2$X-HTL.2** after lyophilization as a white solid.

**$^1$H NMR** (600 MHz, MeOD-d4) δ 9.19 (t, *J* = 5.9 Hz, 1H), 8.21 (dd, *J* = 8.1, 1.3 Hz, 1H), 8.15 (d, *J* = 8.1 Hz, 1H), 7.70 (d, *J* = 8.3 Hz, 2H), 7.68 (s, 1H), 7.38 (d, *J* = 8.2 Hz, 2H), 7.17 (d, *J* = 9.3 Hz, 2H), 7.03 (d, *J* = 7.5 Hz, 1H), 6.95 – 6.91 (m, 4H), 6.72 (s, 1H), 6.60 (dd, *J* = 7.7, 1.3 Hz, 1H), 4.56 (d, *J* = 5.7 Hz, 2H), 3.76 (s, 2H), 3.64 (t, *J* = 6.9 Hz, 2H), 3.59 (t, *J* = 5.6 Hz, 3H), 3.49 (dt, *J* = 16.1, 6.0 Hz, 6H), 2.83 (t, *J* = 6.9 Hz, 2H), 2.68 – 2.63 (m, 2H), 2.00 – 1.95 (m, 2H).
**$^{19}$F NMR** (564 MHz, MeOD-d4) δ -77.4, -110.5 ($m_C$, *J* = 7.4 Hz).
**HRMS** (ESI): calc. for $C_{43}H_{38}ClF_2N_2O_7$ $[M+H]^+$: 767.2330, found: 767.2444.

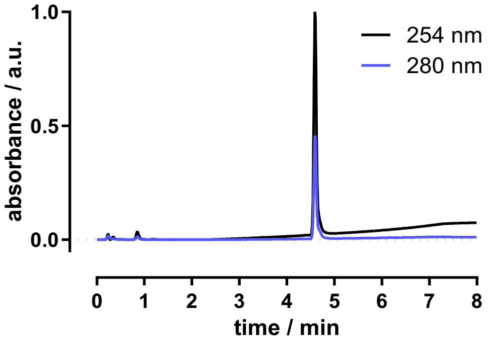

#### 3-(*N*-(2-(4-(3-Chloropropyl)-2-methoxyphenethoxy)ethyl)-3',6'-difluoro-3-oxo-3*H*-spiro[isobenzofuran-1,9'-xanthene]-6-carboxamido)propane-1-sulfonate, F₂X-SHTL.2

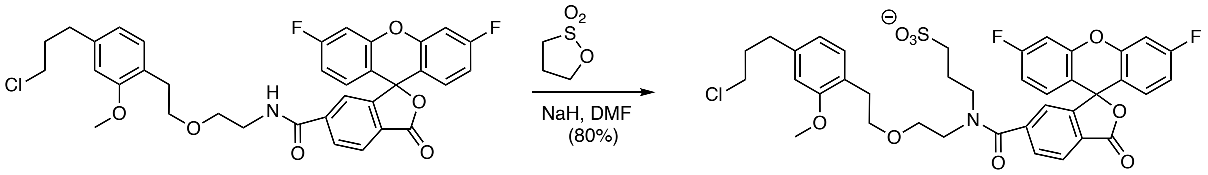

**$F_2$X-preHTL.2** **$F_2$X-SHTL.2**

An Eppendorf tube was charged with **$F_2$X-preHTL.2** (0.5 mg, 0.80 µmol, 1.0 equiv.) in DMF (100 µL). NaH (60% in mineral oil; 0.64 mg, 16 µmol, 20.0 equiv.) and 1,3-propanesultone (0.49 mg, 4.0 µmol, 5.0 equiv.) were added successively. The reaction mixture was incubated at r.t. until complete conversion was observed (by LCMS). The mixture was diluted with $H_2O$:MeCN:AcOH (25:25:1) and subjected to RP-HPLC (MeCN:$H_2O$+0.1% TFA = 50:90 to 95:5 over 40 minutes) to obtain 0.49 mg (80%, TFA salt) of **$F_2$X-SHTL.2** after lyophilization as a white solid.

**$^1$H NMR** (600 MHz, MeOD-d4) δ: 8.03 (d, *J* = 7.9 Hz, 1H), 7.69 (d, *J* = 8.1 Hz, 1H), 7.27 (s, 1H), 7.14 (dd, *J* = 9.4, 2.1 Hz, 2H), 6.97 (d, *J* = 6.9 Hz, 1H), 6.93-6.86 (m, 3H), 6.82 (d, *J* = 7.5 Hz, 1H), 6.68 (s, 1H), 6.59 (d, *J* = 7.5 Hz, 1H), 3.75 (s, 2H), 3.73 (s, 3H), 3.65 (s, 2H), 3.58 (t, *J* = 7.2 Hz, 2H), 3.51 (t, *J* = 6.4 Hz, 2H), 3.36 (d, *J* = 5.2 Hz, 2H), 3.34-3.32 (m, 2H), 2.65 (t, *J* = 7.5 Hz, 2H), 2.47 (t, *J* = 6.7 Hz, 2H), 2.00 (m, 4H).
**$^{19}$F NMR** (564 MHz, MeOD-d4) δ: -76.9, -110.4 ($m_C$, *J* = 7.6 Hz).
**HRMS** (ESI): calc. for $C_{38}H_{35}ClF_2NO_9S^-$ [M+H]$^+$: 756.1846, found: 756.1818.

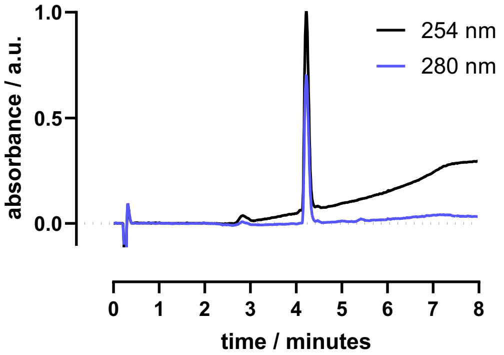

#### 3',6'-Bis(ethylamino)-3-oxo-3*H*-spiro[isobenzofuran-1,9'-xanthene]-6-carboxylic acid, (EtNH)$_2$X-6-COOH

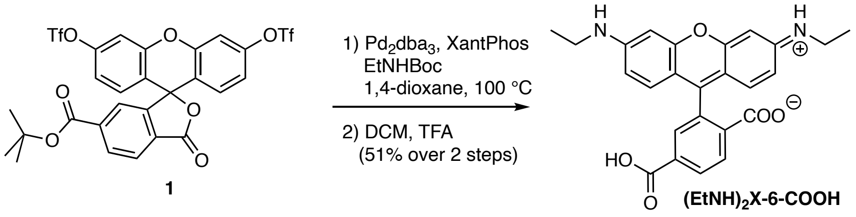

A Schlenk tube was charged with 6-*tert*-butoxycarbonylfluorescein ditriflate **1** (10.0 mg, 14.4 µmol), $Pd_2dba_3$ (2.0 mg, 2.18 µmol, 0.15 equiv.), XPhos (2.1 mg, 4.41 µmol, 0.30 equiv.), $Cs_2CO_3$ (23.4 mg, 71.8 µmol, 5 equiv.), *tert*-butyl ethylcarbamate (10.0 mg, 68.9 µmol, 4.8 equiv.) were added and dissolved with *dry* dioxane (250 µL) under $N_2$. The Schlenk tube was sealed and evacuated/backfilled with $N_2$ (3×). The reaction mixture was stirred at 100 °C over 14 h. After cooling to r.t., the crude mixture was filtrated through a plug of Celite and washed with MeOH than concentrated under vacuum. The residue obtained was dissolved in a mixture of DCM/TFA (3:1) during 3h at rt. The reaction was concentrated under vacuum and dissolved in DMSO:$H_2O$:MeCN:AcOH (10:25:25:1) and subjected to RP-HPLC (MeCN:$H_2O$+0.1% TFA = 10:90 to 90:10 over 60 minutes), to afford 4.6 mg (51%, TFA salt) of **3** as an orange solid.

**$^1$H NMR** (600 MHz, MeOD-d4) δ 8.41 – 8.36 (m, 2H), 7.96 (s, 1H), 7.07 – 6.99 (m, 2H), 6.88 – 6.78 (m, 4H), 3.42 (q, $J$ = 7.3 Hz, 4H), 1.34 (t, $J$ = 7.2 Hz, 6H).

**$^{13}$C NMR** (151 MHz, MeOD-d4) δ 167.9 (C), 167.7 (C), 160.1 (C), 159.3 (C), 136.5 (C), 135.9 (CH), 135.4 (CH), 132.7 (2 × CH), 132.2 (2 × CH), 115.0 (C), 39.1 (N-$CH_2$), 14.0 ($CH_3$). (One Cq is missing).

**$^{19}$F NMR** (564 MHz, MeOD-d4) δ -77.0.

**HRMS** (ESI): calc. for $C_{25}H_{23}N_2O_5$ $[M+H]^+$: 431.1602, found: 431.1623.

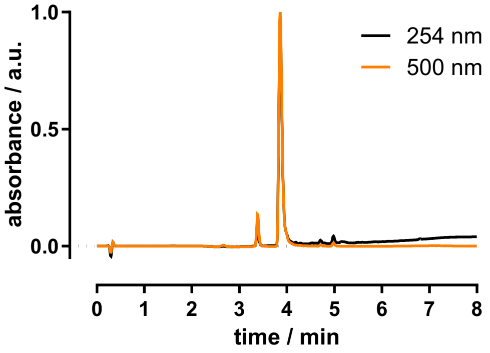

Remark: Rhodol derivative 1.0 mg (17%, TFA salt) was also isolated as an orange solid.

#### *N*-(2-(4-(3-Chloropropyl)-2-methoxyphenethoxy)ethyl)-3',6'-bis(ethylamino)-3-oxo-3H-spiro[isobenzofuran-1,9'-xanthene]-6-carboxamide, (EtNH)$_2$X-preHTL.2

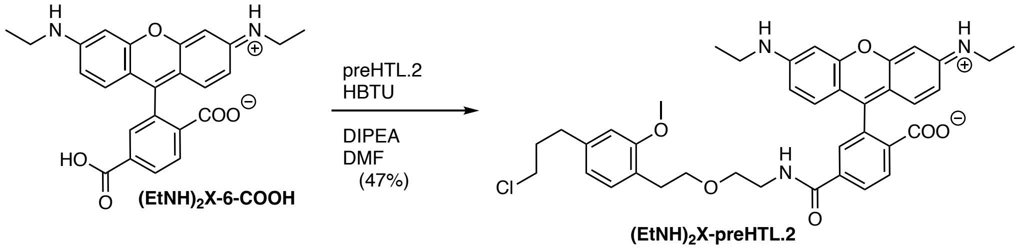

An Eppendorf tube was charged with **(EtNH)$_2$X-6-COOH** (0.8 mg, 1.90 µmol, 1.0 equiv.) in DMF (100 µL). DIPEA (2.5 µL, 15 µmol, 8.0 equiv.) and a 100 mM solution of TSTU (28 µL, 1.70 µmol, 1.5 equiv.) in DMF were added successively. The reaction mixture was allowed to incubate for 15 minutes, and then a 100 mM HTL-2.0-NH$_2$ solution (28 µL, 1.70 µmol, 1.5 equiv.) was added in one drop. The reaction mixture was incubated at r.t. until complete conversion was observed (by LCMS). The mixture was diluted with $H_2O$:MeCN:AcOH (25:25:1) and subjected to RP-HPLC (MeCN:$H_2O$+0.1% TFA = 10:90 to 90:10 over 60 minutes) to obtain 0.7 mg (47%, TFA salt) of **(EtNH)$_2$X-preHTL.2** after lyophilization as an orange solid.

**$^1$H NMR** (600 MHz, MeOD-d4) δ 8.66 (t, *J* = 5.5 Hz, 1H), 8.39 (d, *J* = 8.2 Hz, 1H), 8.15 (dd, *J* = 8.2, 1.7 Hz, 1H), 7.79 (d, *J* = 1.7 Hz, 1H), 7.04 – 6.99 (m, 2H), 6.85 – 6.82 (m, 4H), 6.74 (s, 1H), 6.60 (dd, *J* = 7.5, 1.2 Hz, 1H), 3.77 (s, 3H), 3.63 (q, *J* = 6.2 Hz, 4H), 3.56 (q, *J* = 5.3 Hz, 2H), 3.51 (t, *J* = 6.5 Hz, 2H), 3.42 (q, *J* = 7.2 Hz, 4H), 2.81 (t, *J* = 7.0 Hz, 2H), 2.71 – 2.66 (m, 2H), 2.01 (q, *J* = 6.6 Hz, 2H), 1.34 (t, *J* = 7.2 Hz, 6H).
**$^{19}$F NMR** (564 MHz, MeOD-d4) δ -77.0.
**HRMS** (ESI): calc. for $C_{39}H_{43}ClN_2O_6$ $[M+H]^+$: 684.2835, found: 684.2874.

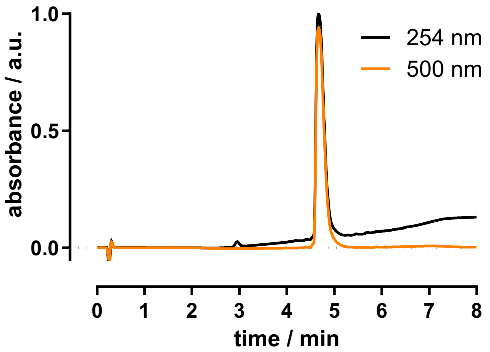

### NMR analysis

#### $^{1}$H NMR (600 MHz, MeOD-d4) of F$_2$X-6-COOH

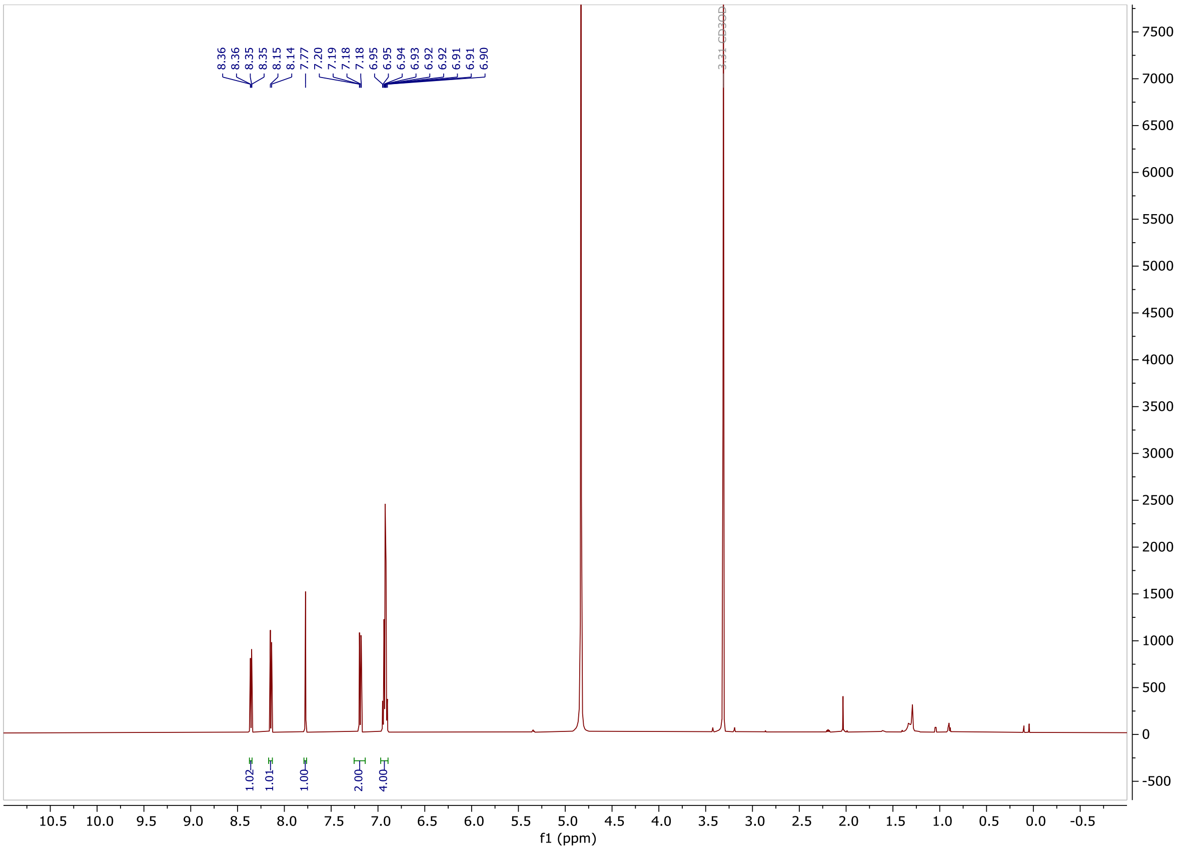

#### $^{13}$C NMR (151 MHz, MeOD-d4) of F$_2$X-6-COOH

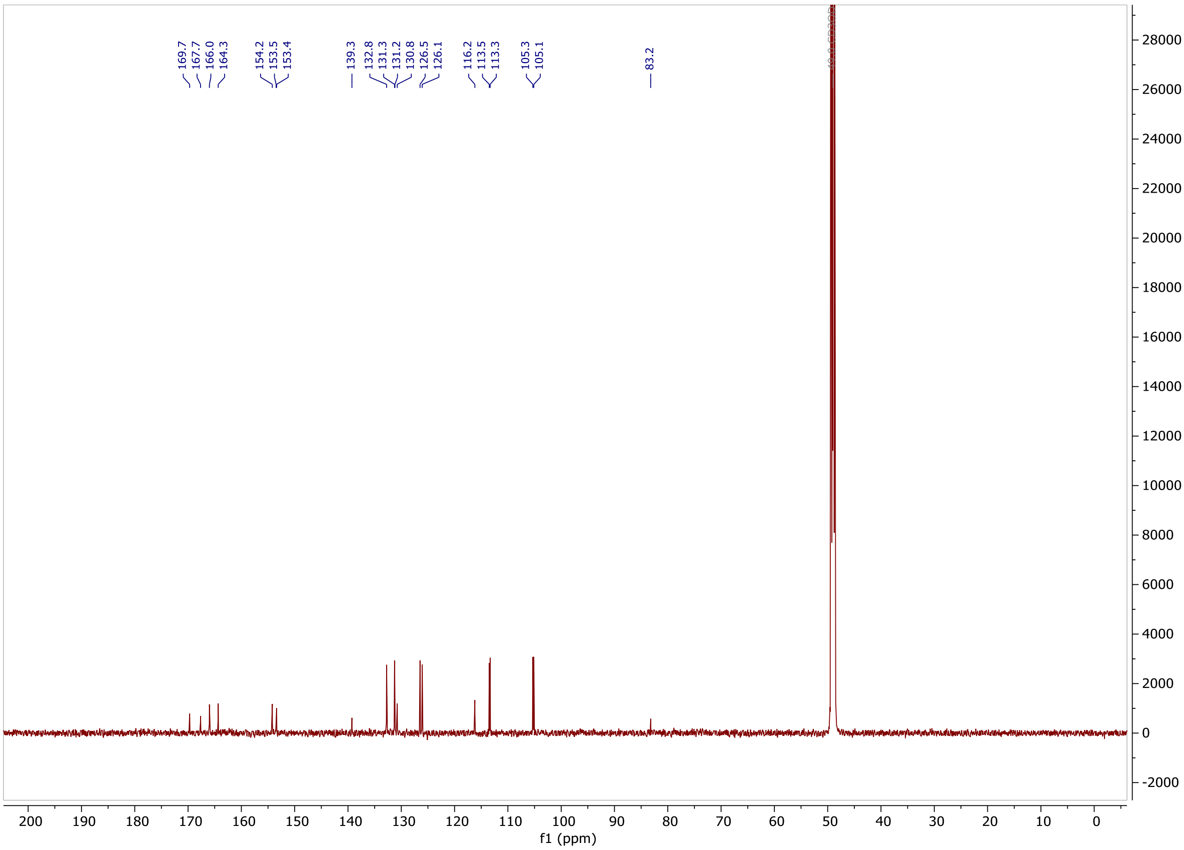

**$^{19}$F NMR** (564 MHz, MeOD-d4) of **$F_2$X-6-COOH**

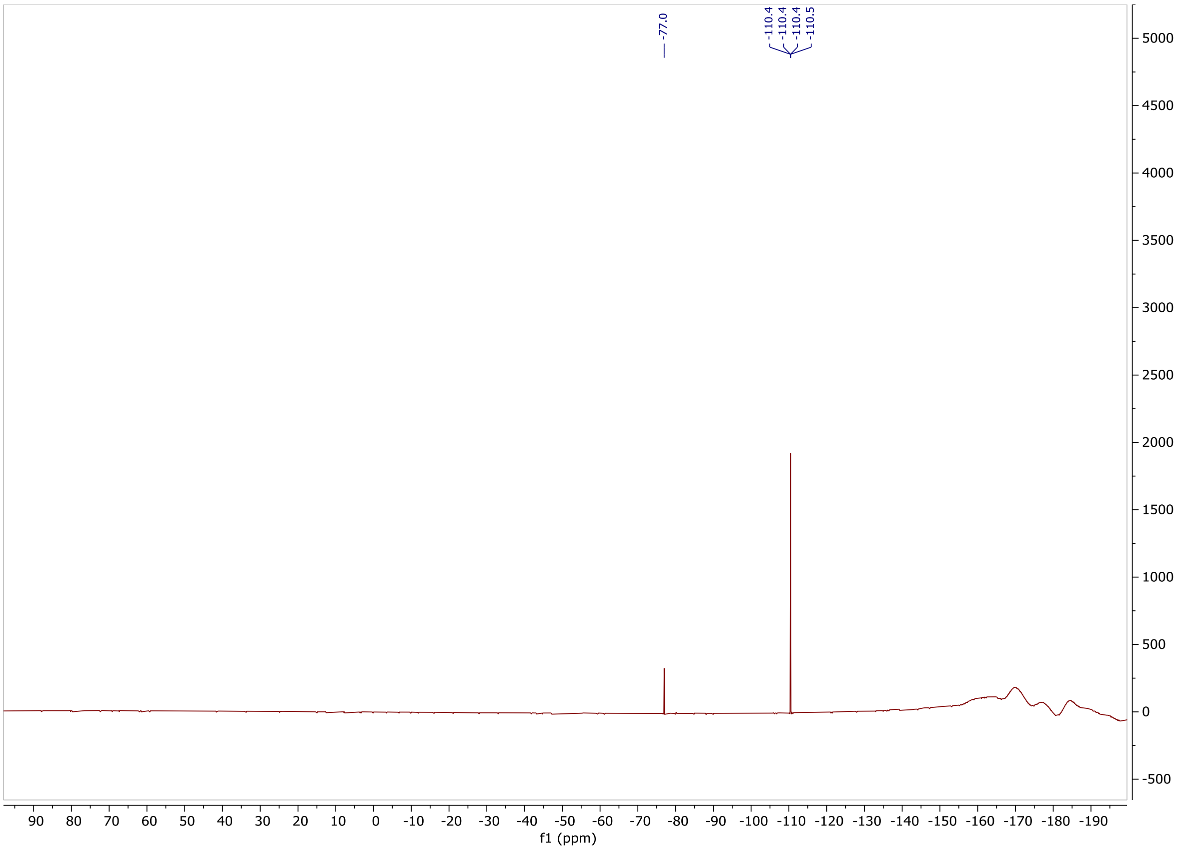

**$^{1}$H NMR** (600 MHz, MeOD-d4) of **$F_2$X-HTL.1**

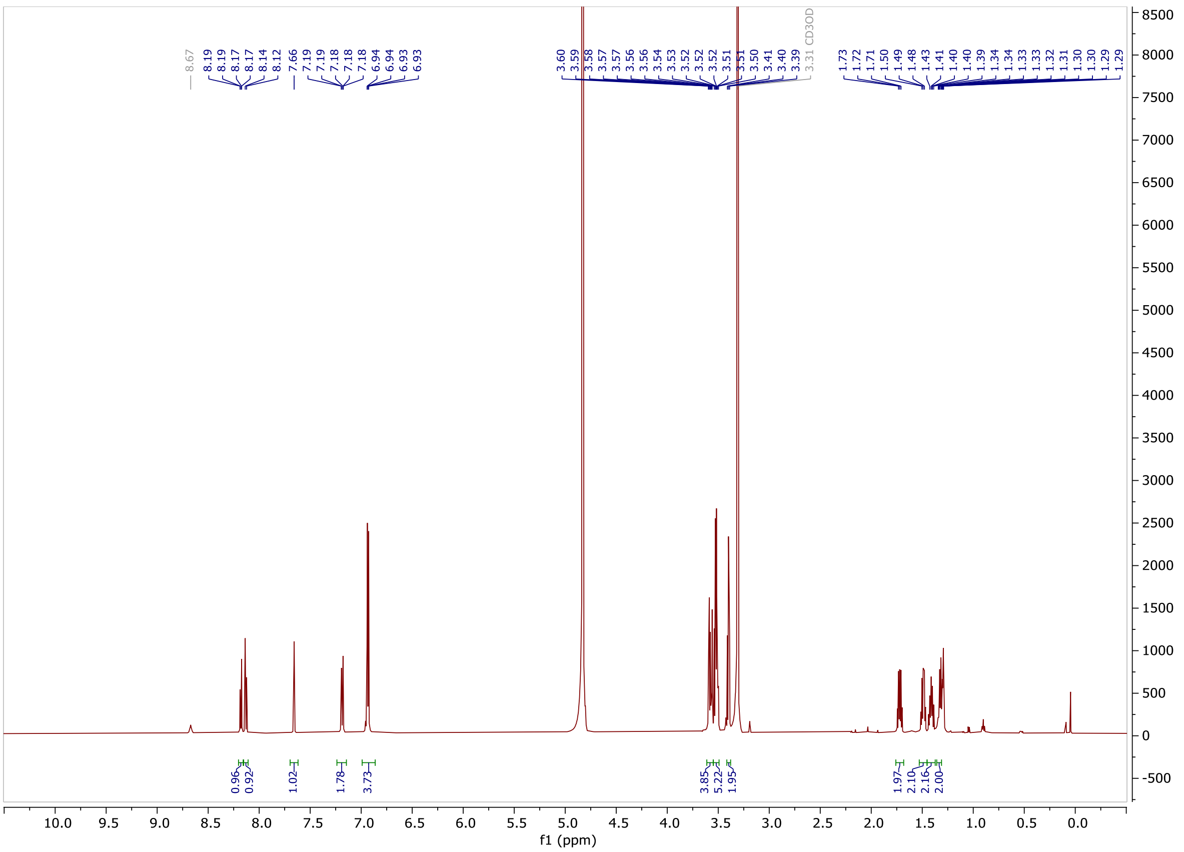

**$^{19}$F NMR** (564 MHz, MeOD-d4) of **$F_2$X-HTL.1**

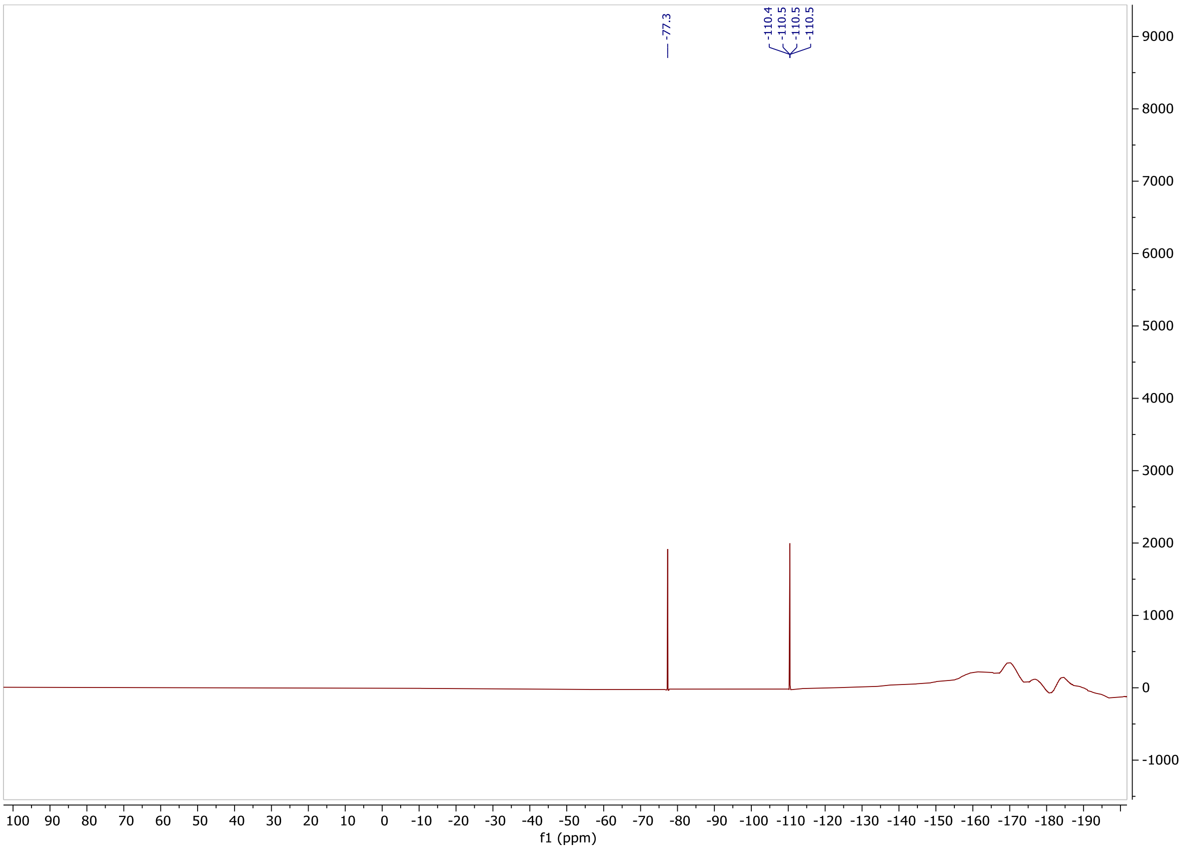

$^{1}$H **NMR** (600 MHz, MeOD-d4) of **$F_2$X-preHTL.2**

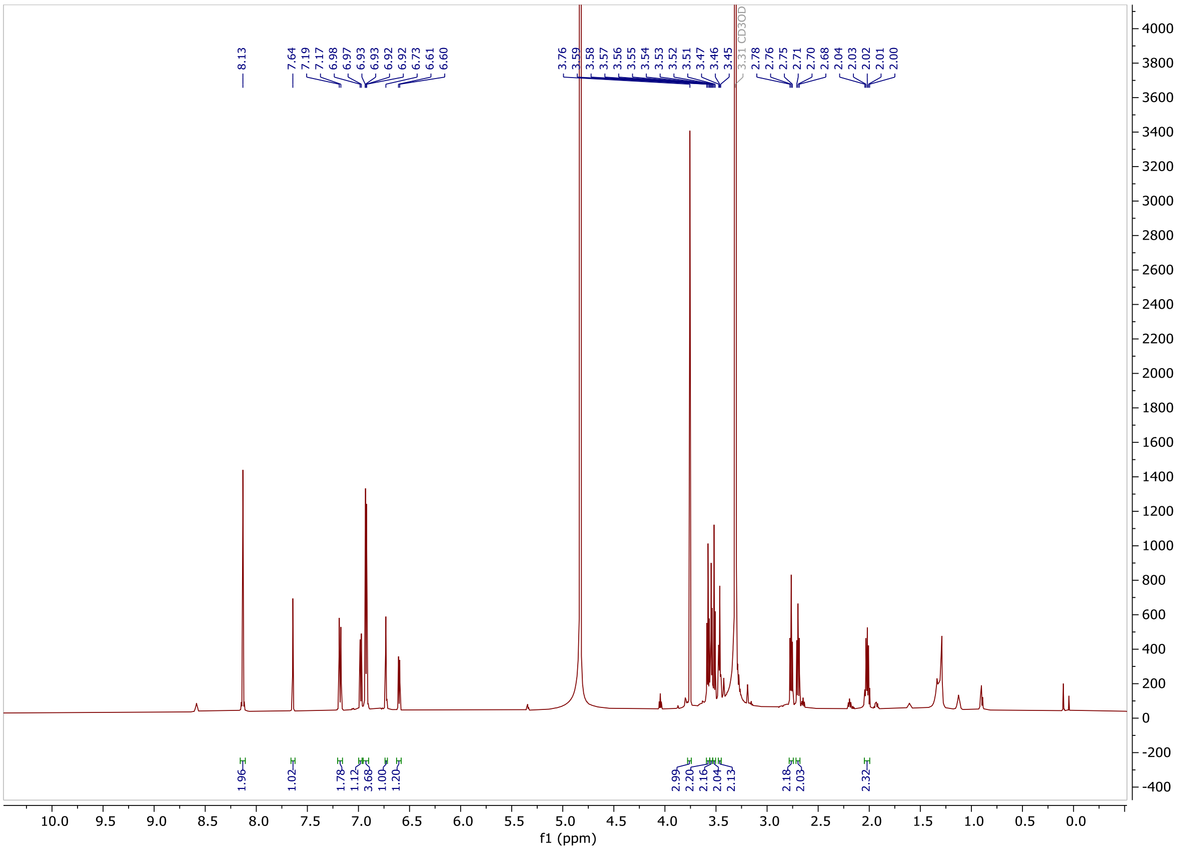

$^{13}$C **NMR** (151 MHz, MeOD-d4) of **$F_2$X-preHTL.2**

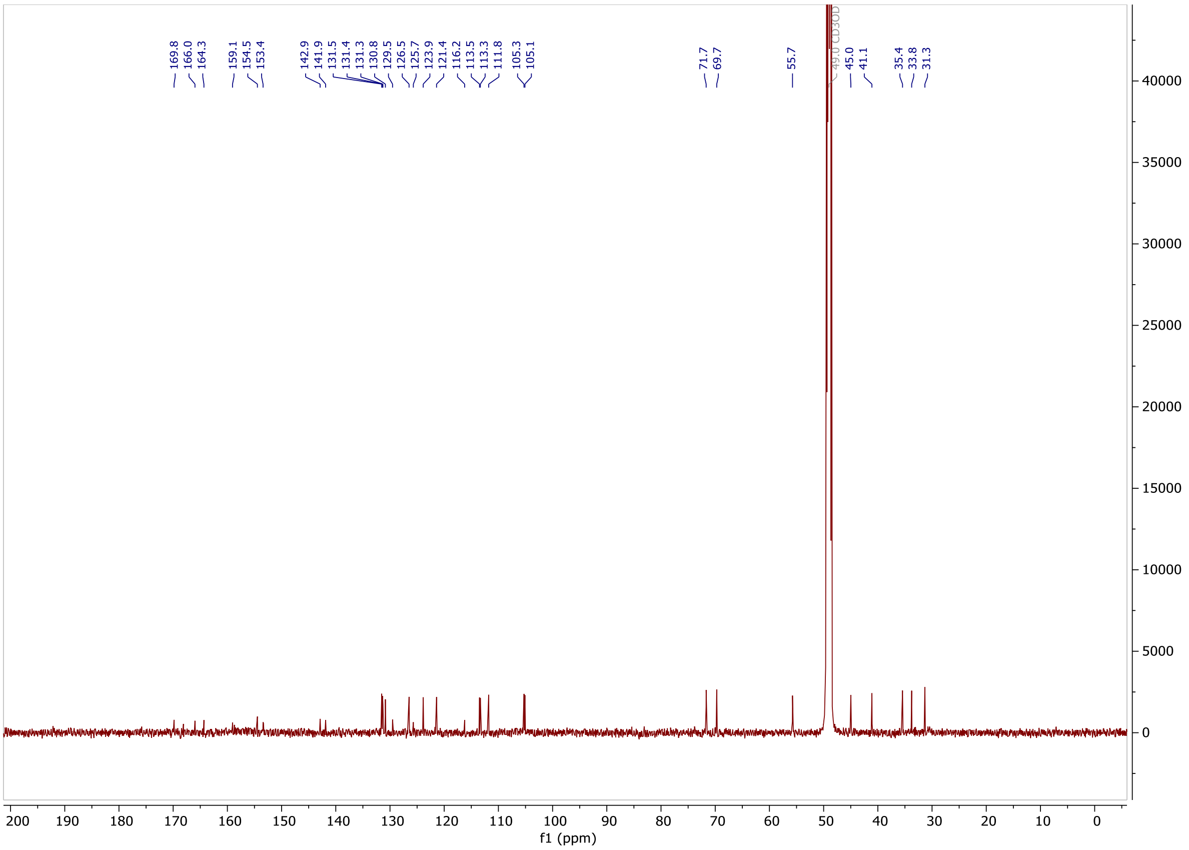

**[19]F NMR** (564 MHz, MeOD-d4) of **$F_2$X-preHTL.2**

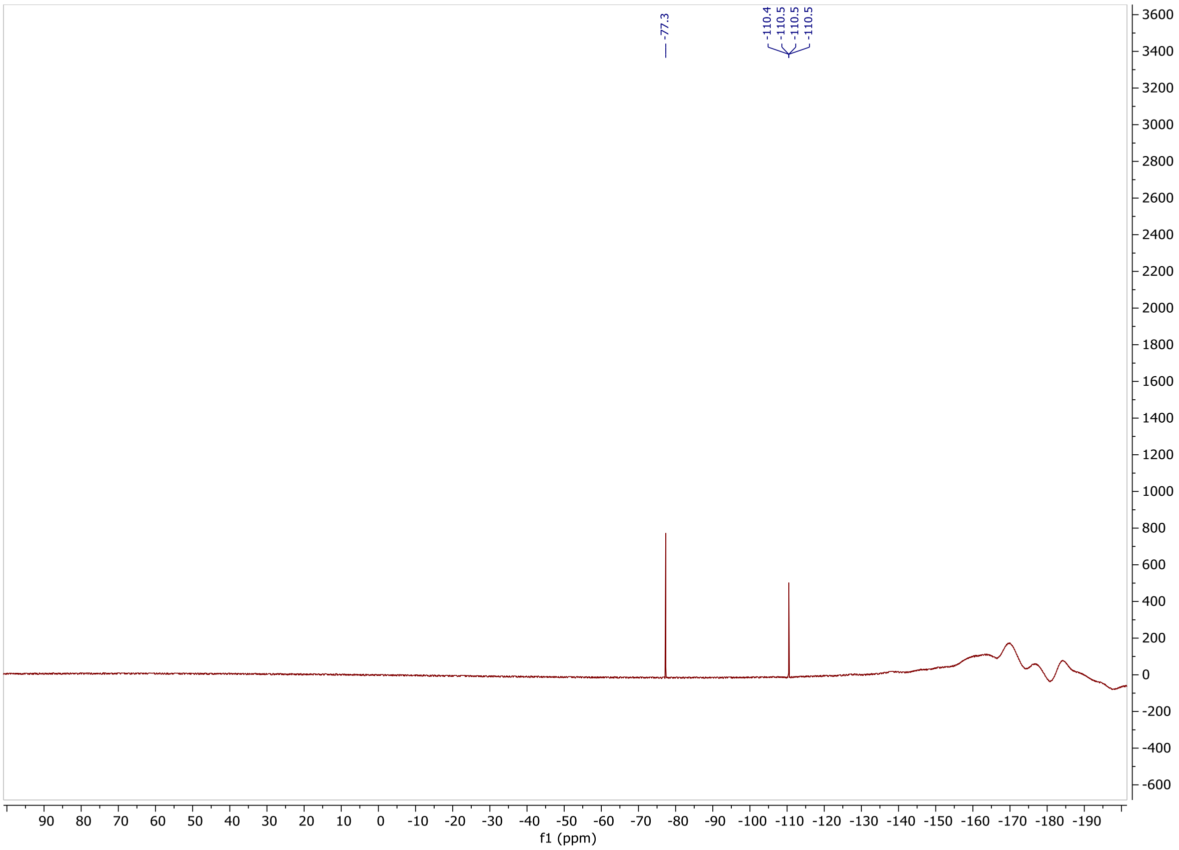

[1]H NMR (600 MHz, MeOD-d4) of $F_2$X-HTL.2

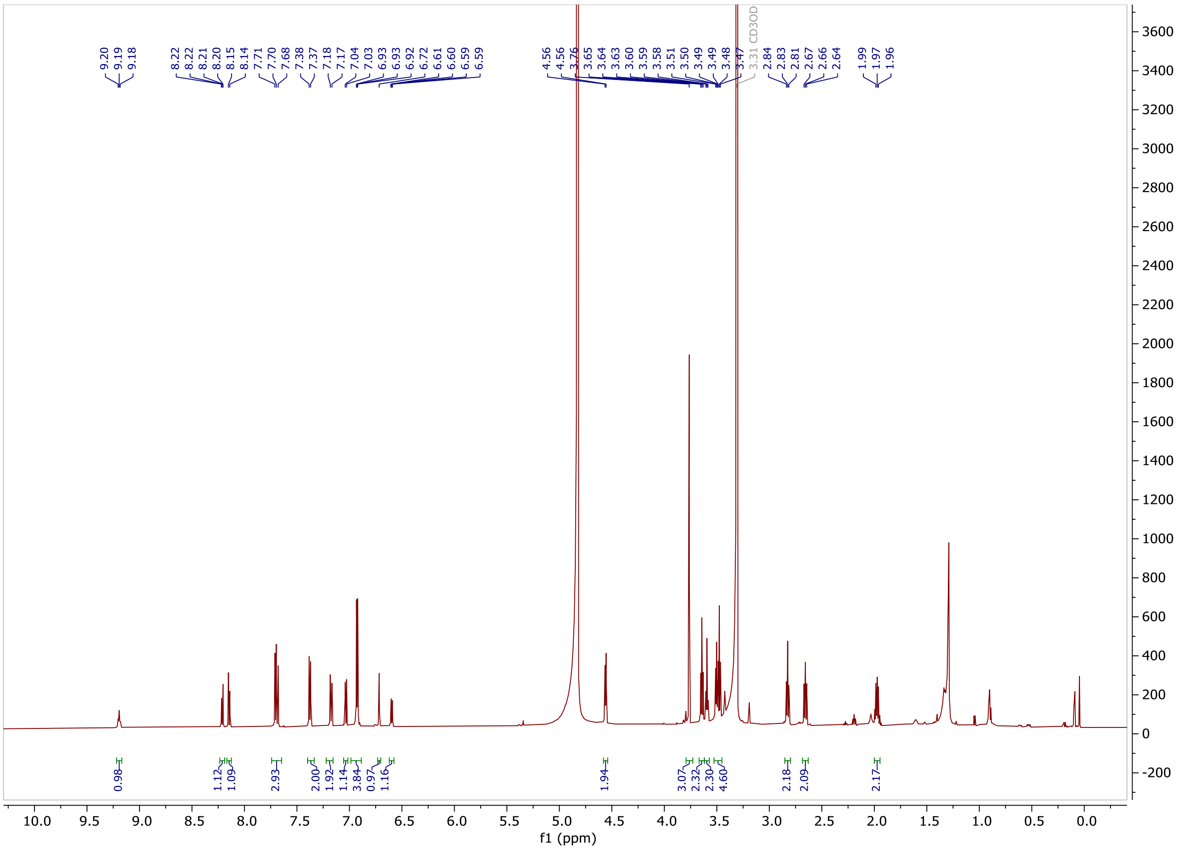

[19]F NMR (564 MHz, MeOD-d4) of $F_2$X-HTL.2

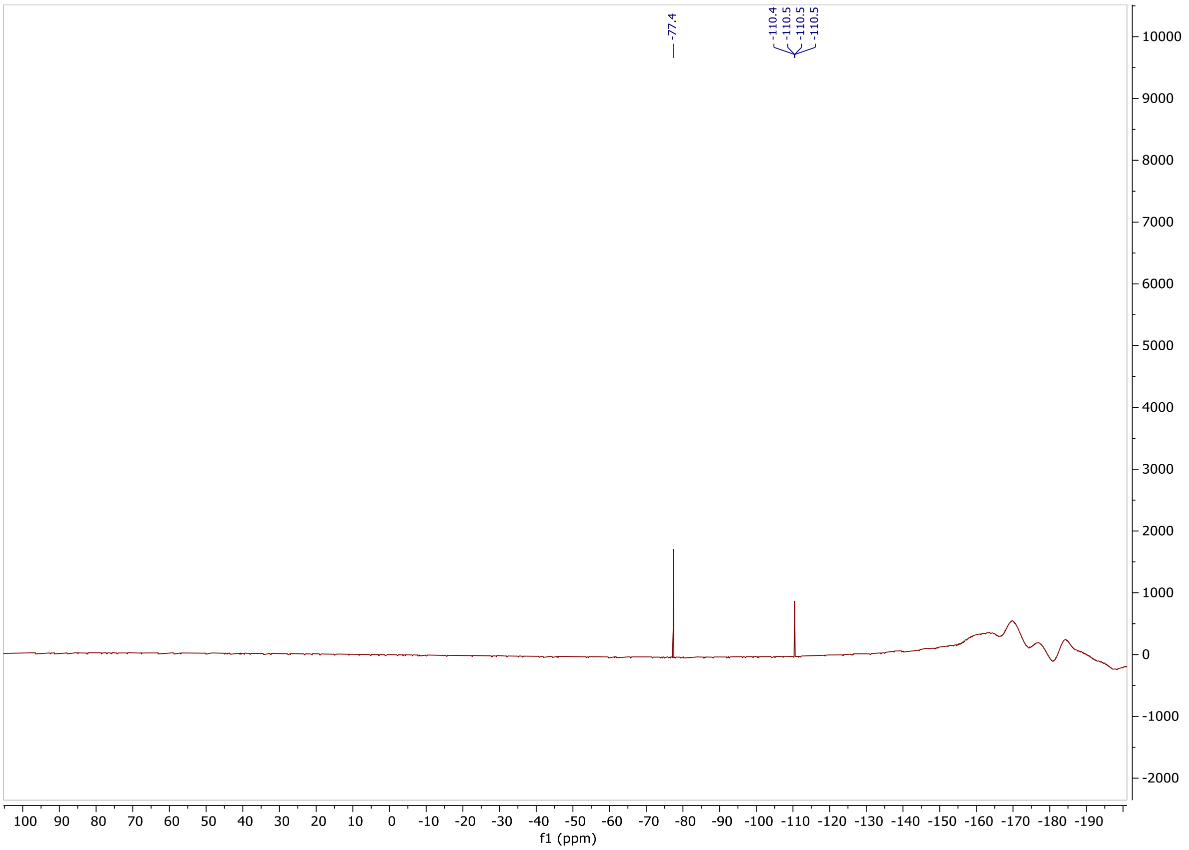

**$^{1}$H NMR** (600 MHz, MeOD-d4) of **$F_2$X-SHTL.2**

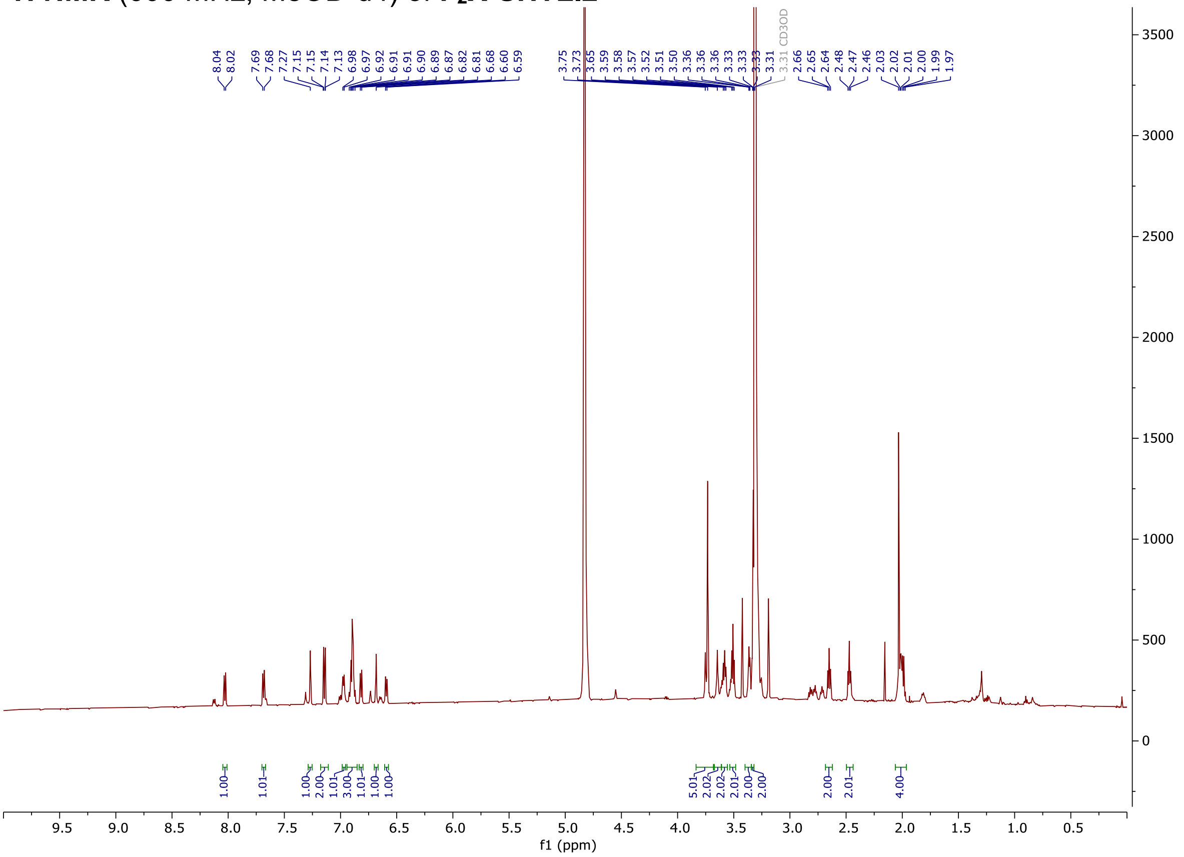

**$^{19}$F NMR** (564 MHz, MeOD-d4) of of **$F_2$X-SHTL.2**

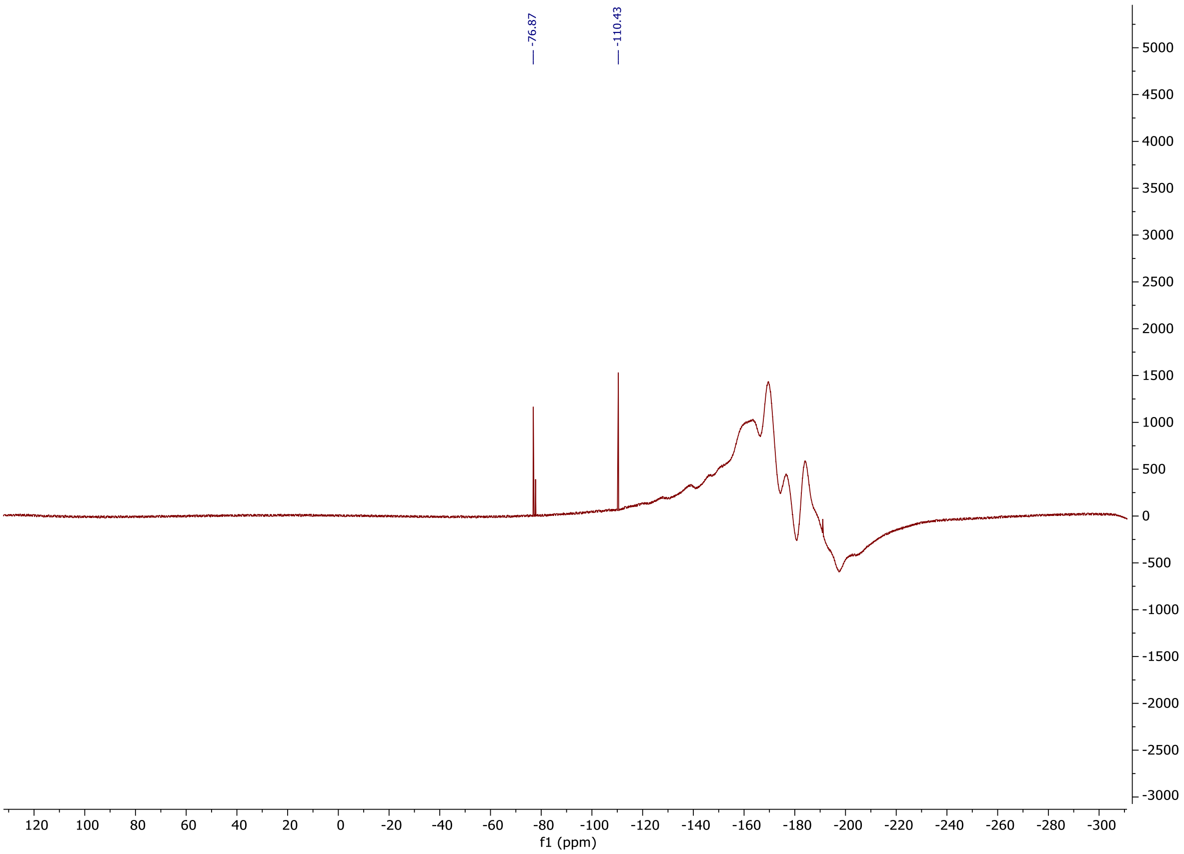

**$^{1}$H NMR** (600 MHz, MeOD-d4) of **(EtNH)$_2$X-6-COOH**

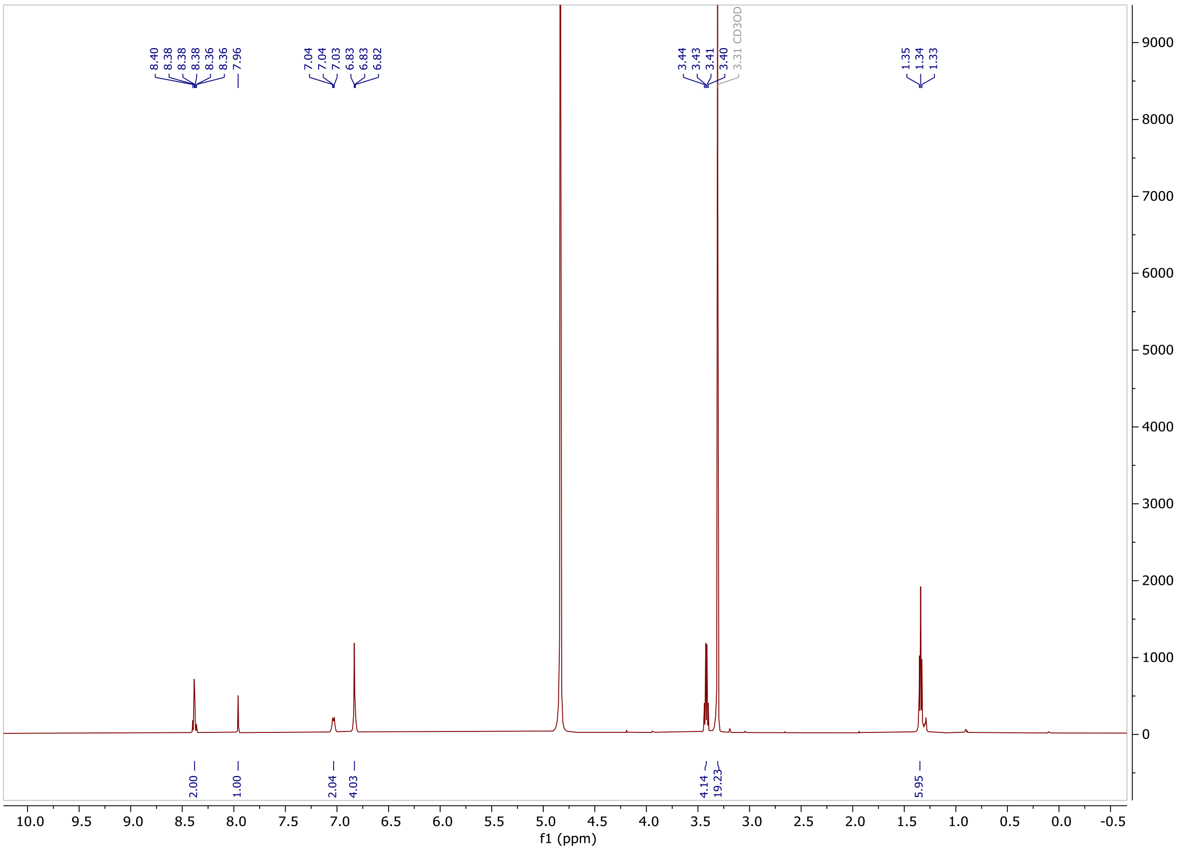

**$^{13}$C NMR** (151 MHz, MeOD-d4) of **$(EtNH)_2$X-6-COOH**

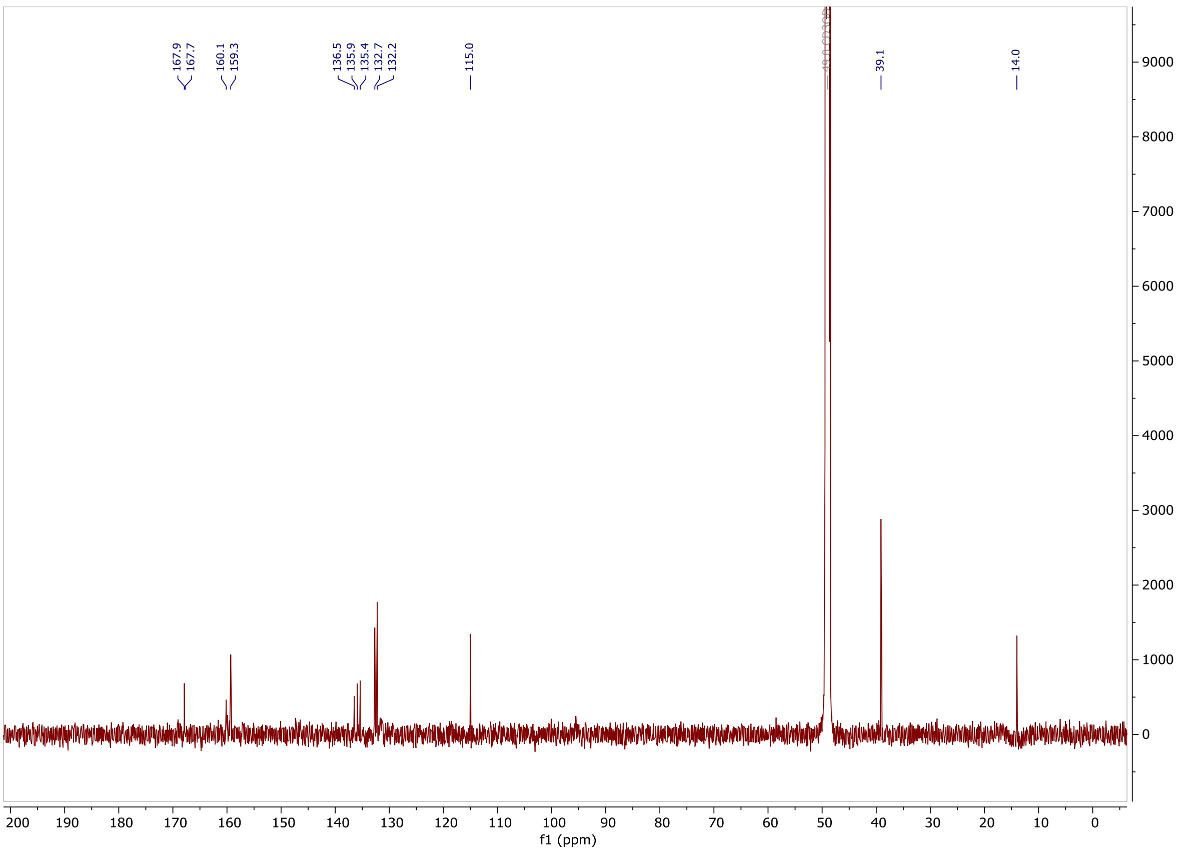

**$^{19}$F NMR** (564 MHz, MeOD-d4) of **(EtNH)$_2$X-6-COOH**

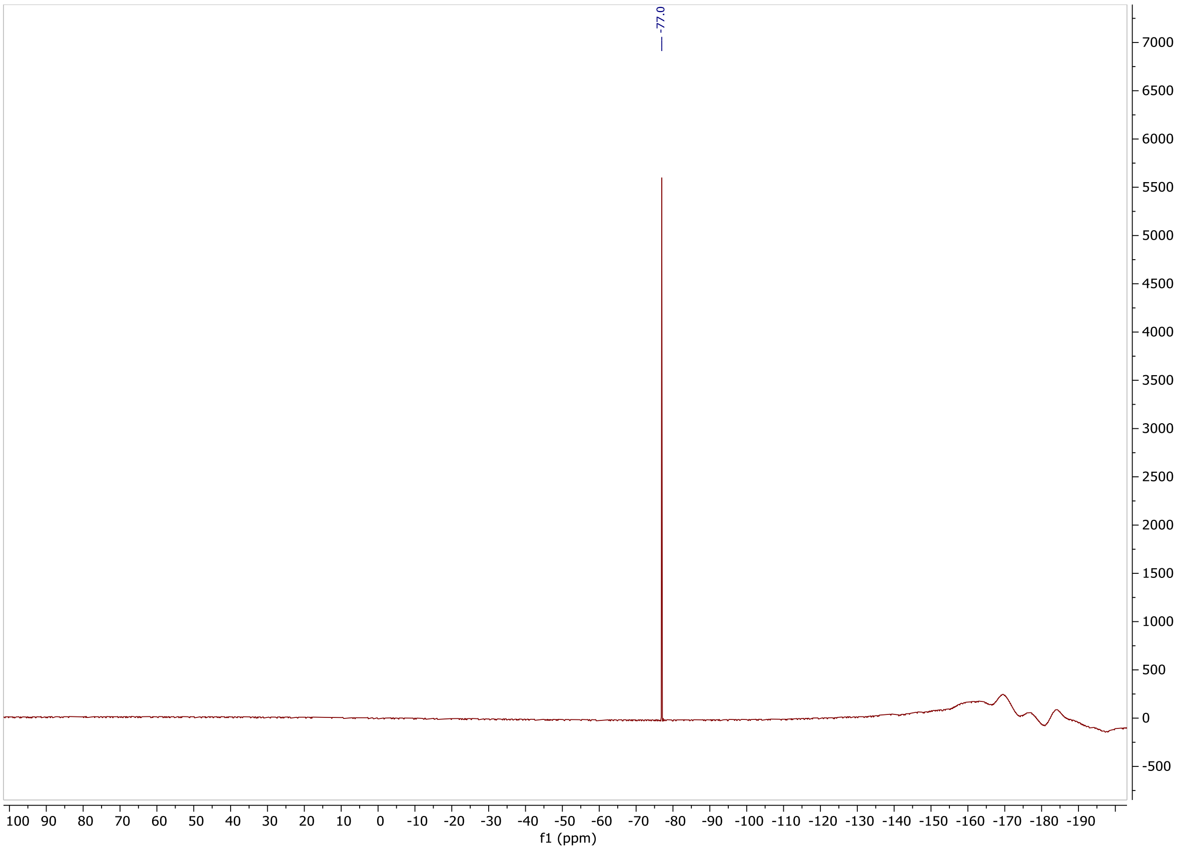

**$^{1}$H NMR** (600 MHz, MeOD-d4) of **(EtNH)$_2$X-preHTL.2**

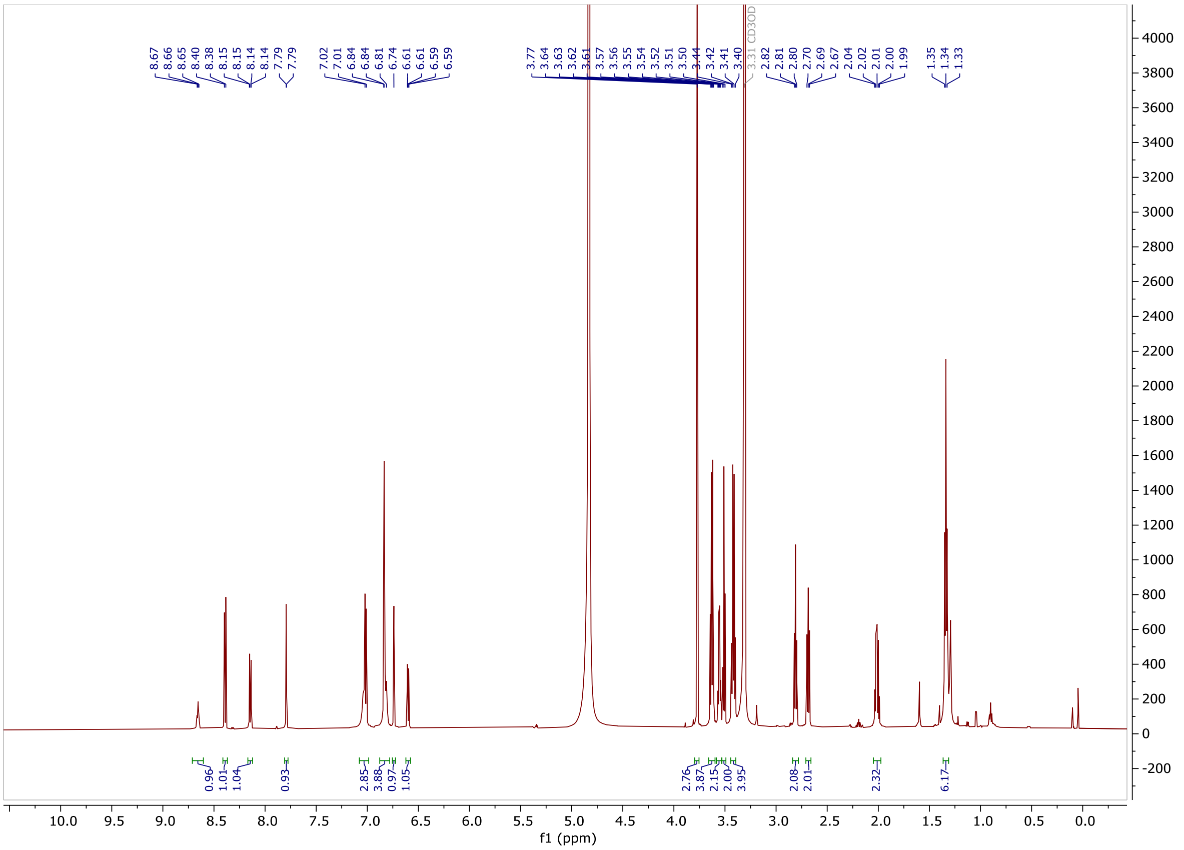

**$^{19}$F NMR** (564 MHz, MeOD-d4) of **(EtNH)$_2$X-preHTL.2**

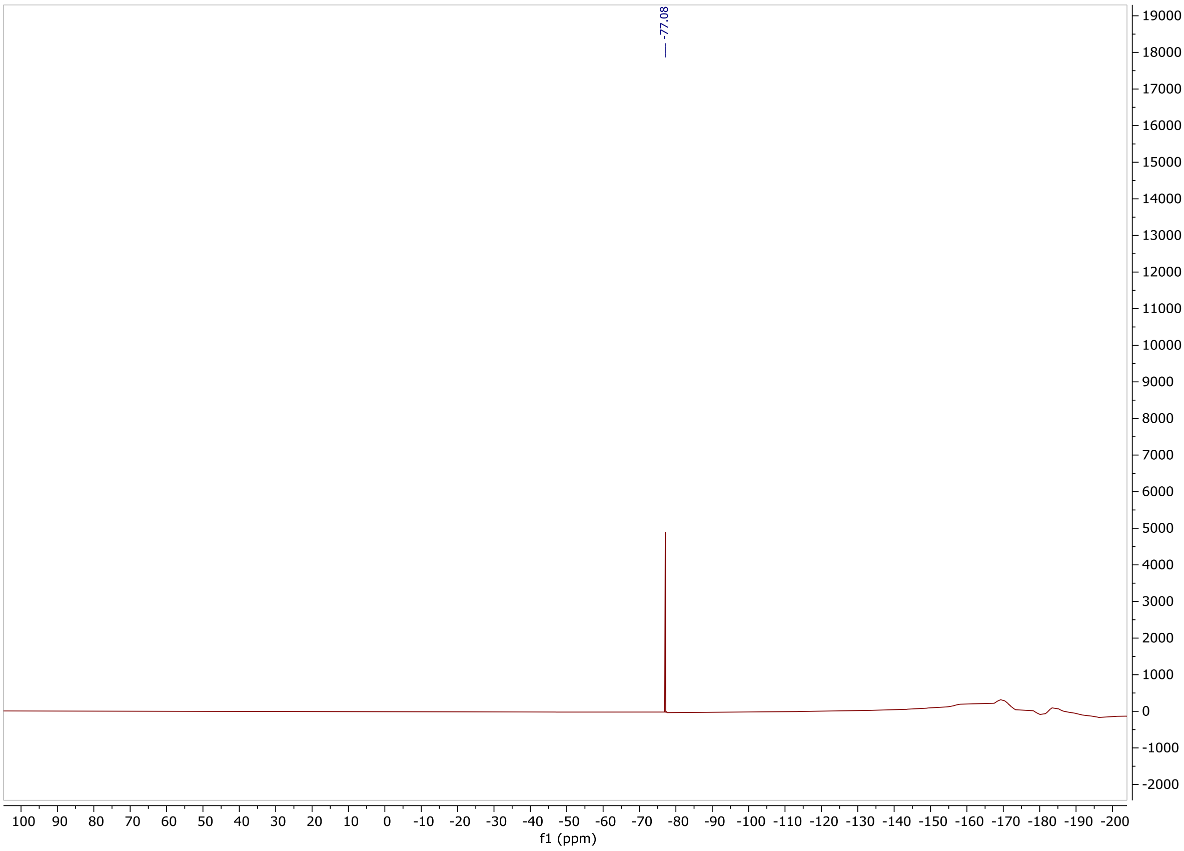
